# Predicting how perturbations reshape cellular trajectories with PerturbGen

**DOI:** 10.64898/2026.03.04.709254

**Authors:** Kevin Chi Hao Ly, Adib Miraki Feriz, Tomoya Isobe, Amirhossein Vahidi, Delshad Vaghari, Anthony Rostron, Mariana Quiroga Londoño, Nicole Mende, MS Vijayabaskar, Marie Moullet, April Rose Foster, Emily Graves, Fereshteh Torabi, Chloe Admane, David Horsfall, Daniela Basurto-Lozada, Emily Stephenson, Rachel A. Botting, Deena Iskander, Simone Webb, Issac Goh, Daniyal J. Jafree, Katherine H. M. Sturgess, Jonathan Scott, Wezi Sendama, Laura Magnani, Rebecca Hannah, Hesam Asadollahzadeh, Ash Holland, Martin Prete, John Simpson, Anindita Roy, Irene Roberts, Laura Jardine, Elisa Laurenti, Gosia Trynka, Nicola K. Wilson, Muzlifah Haniffa, Berthold Göttgens, Mo Lotfollahi

## Abstract

A major challenge in biology is predicting how cells transition between states over time and how perturbations disrupt these transitions. Understanding such dynamics is critical for identifying interventions that reverse pathological programs or reprogram cells toward desired states. Although recent computational approaches can predict single-cell perturbation responses *in silico*, they cannot predict responses across dynamic cell trajectories, for example how early perturbations reconfigure later cell states. To address this gap, we introduce PerturbGen, a generative foundation model trained on over 100 million single-cell transcriptomes that predicts perturbation responses along cellular trajectories. PerturbGen predicts how genetic perturbation at source state shapes downstream states, alters gene programs and trajectories across time, for example in differentiation or disease progression. We apply PerturbGen to three newly generated multi-condition human single-cell datasets spanning immune responses, hematopoiesis and skin development. In an *in vivo* immune challenge, PerturbGen predicts that knocking out an *IL1B* signal in myeloid cells attenuates later cytokine-interferon programs, with downstream changes consistent with a reversal of IL-1β stimulation signature. In hematopoiesis, anchoring perturbation-induced programs to human genetics enables simulation of monogenic blood disorders, recapitulates established disease-associated biology whilst systematically revealing lineage-specific programs, including in lineages where this was not previously possible. In skin organoids, PerturbGen predicted that Wnt activation enhances stromal differentiation recapitulating the trajectory observed in human prenatal skin, findings that were functionally validated by experimentally activating Wnt signaling. Together, PerturbGen extends modeling of gene perturbations from static to dynamic cellular systems. We envision PerturbGen enabling the creation of *in silico*, trajectory-aware perturbation atlases and virtual cells across diverse biological scenarios, supporting optimization of disease models and prioritization of candidate molecular interventions for therapeutic discovery.

## Introduction

Cellular identity and function are not static properties but emerge from transitions between states that unfold across time. Transitions in cells from one state to another, defined as trajectories, are a fundamental property of biological tissues across diverse contexts. Examples of cell state transitions include the progression of cells along differentiation trajectories during development^1,2^ and the constant replenishment of organs with rapid cell turnover, such as skin, gut and blood^3–7^. Genetic, molecular or chemical perturbations, when introduced at specific points along a trajectory, can redirect cell state, alter homeostasis and function, or give rise to pathological states^8^. Understanding how cellular states evolve in development and disease, and how perturbations reshape these trajectories, is central to profiling developmental biology, dissecting disease mechanisms and informing therapeutic interventions.

Despite its importance, systematically exploring how perturbations reshape cellular trajectories remains experimentally challenging. Testing perturbations across all cellular states and time points, particularly in combinatorial designs, is rarely feasible at scale. Although pooled CRISPR-based technologies^9,10^ enable genome-scale perturbation at single-cell resolution, their applicability to *in vivo* systems and their ability to temporally resolve trajectories remain limited. Efforts to perform physiologically-relevant tissue perturbation experiments often depend on complex culture systems^11,12^ that are difficult to establish and incompletely capture the diversity of cellular and developmental states present *in vivo*^13^. Together, these constraints underscore the need for scalable computational frameworks for assessing perturbation effects across cellular trajectories.

Computational approaches for modeling cellular perturbations have advanced substantially in recent years, encompassing intrinsic perturbations such as genetic changes or extrinsic perturbations such as chemical or molecular stimuli. Existing methods primarily predict responses of previously unseen perturbations^14–16^ at single-cell resolution, including with task-specific neural networks^17–22^, fine-tuning or adapting single-cell foundation models^23–29^, and gene regulatory network (GRN)-based frameworks that propagate perturbations through inferred regulatory interactions. However, for approaches that depend on inferred regulatory networks, the scope of possible perturbations is constrained by network coverage and confidence, limiting systematic interrogation of perturbation effects across the transcriptome and cellular states. Moreover, perturbation prediction is typically formulated within a single observed cellular state in a fixed context, often derived from immortalized or *ex vivo* systems. Consequently, existing approaches do not account for how state-dependent perturbation effects along cellular trajectories reshape downstream consequences such as lineage specification or immune activation.

In parallel, methods that model cellular dynamics from observational data^30^, including pseudotime inference^2,31,32^ and RNA velocity^33,34^, can be applied to single-cell RNA sequencing (scRNA-seq) datasets that aim to capture dynamic cellular processes^35^. More recently, generative frameworks based on neural ordinary differential equations^36,37^, optimal transport^38–40^, flow matching^41–43^, or diffusion processes^44^ have been developed to model and predict cellular states at unobserved time points. While these approaches capture cellular trajectories, they are primarily designed to describe unperturbed systems and do not incorporate genetic perturbations. Consequently, existing methods either predict perturbation effects within fixed cellular contexts, or model changes in cellular state without perturbations, leaving a critical gap for approaches that integrate modeling of cell state transitions with perturbation effects.

To address this gap, we introduce PerturbGen, a generative foundation model that predicts state-dependent perturbation responses along cellular trajectories, defined by experimentally observed cell states from stage-resolved single-cell measurements or differentiation-associated developmental stages. PerturbGen enables inference of how perturbations applied at defined trajectory positions influence downstream transcriptional programs and cell fate decisions. We apply PerturbGen to three newly generated human single-cell datasets, identifying interventions that predict downstream inflammatory states in an *in vivo* immune activation challenge, and constructing large-scale virtual perturbation atlases across *in vivo* blood and *in vitro* skin development that provide a comprehensive strategy to map the effects of genetic interventions across cellular trajectories. By integrating predicted perturbation-induced programs (PIPs) with human genetic evidence, alongside targeted experimental evaluation, we establish a framework for exploring perturbation-driven remodelling of cell state transitions during human development, homeostasis and disease. PerturbGen represents a step toward AI-driven construction of virtual cells^45–47^ and *in silico* perturbation atlases that guide experimental design by identifying when and in which cell states interventions are predicted to influence downstream outcomes, enabling more realistic modeling by incorporating dynamic, context-dependent cell and tissue processes that have been underrepresented in perturbation modeling.

## Results

### Generative modeling of cellular trajectories and their response to *in silico* genetic perturbations

PerturbGen models cellular trajectories by predicting gene expression of a target cell state conditioned on a source state and intermediate contextual states along the same trajectory (Fig. 1a; see Methods). Discrete states correspond to experimentally observed conditions, such as time-resolved measurements or developmental and differentiation stages. We define a source state (𝑆₀), a set of contextual states (𝑆_1_,.., 𝑆 ^𝑆′−1^) that precede the target and a target state (𝑆_𝑆′_), which represents a downstream state of interest. For example, along a differentiation trajectory, 𝑆₀ may correspond to a progenitor state, the contextual states to progressively differentiated states, and 𝑆_𝑆′_ to a terminally differentiated cell type. The gene expression profile of each cell is represented as a ranked sequence of gene tokens ordered by normalized expression^24^ (see Methods). Given a tokenized source and context states, PerturbGen generates a token sequence representing the target state using an encoder–decoder architecture^48^, which is subsequently converted back into gene expression. By conditioning target-state generation on both the source and contextual states, PerturbGen explicitly models transitions between states along a trajectory, rather than treating each cellular snapshot independently. This design enables the prediction of downstream effects of *in silico* genetic perturbations by modifying the source state representation while preserving the cellular context. The model then generates predicted gene expression at later stages along the trajectory, allowing direct comparison between perturbed and unperturbed predictions to yield cell state–specific perturbation effects. Scaling this procedure across genes enables systematic *in silico* screening and construction of perturbation atlases spanning intermediate and downstream states (see Methods) and can, in principle, be extended genome-wide.

**Fig 1.**
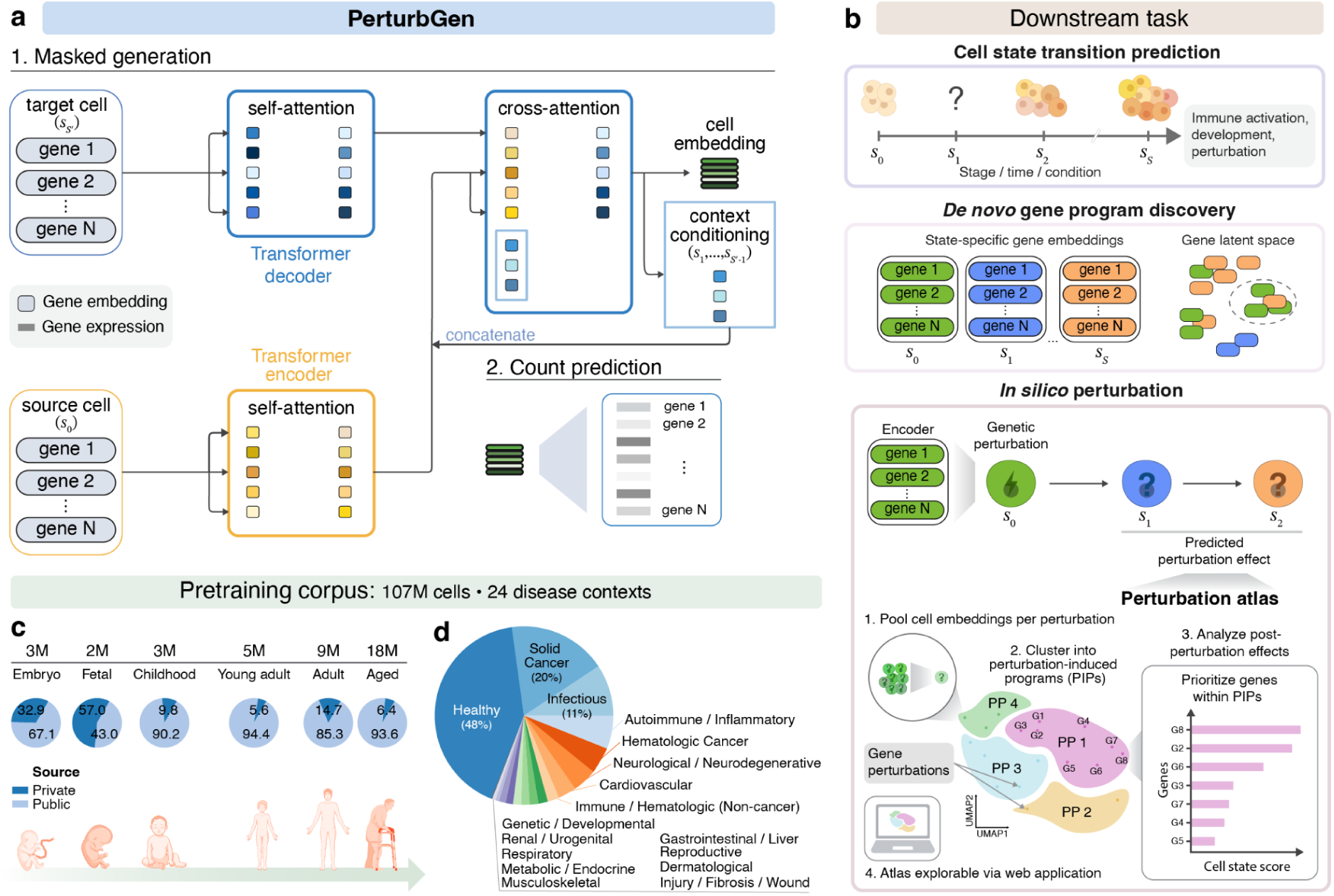
PerturbGen enables generative modeling of cellular trajectories and their response to *in silico* genetic perturbation. **a,** PerturbGen models state-to-state transitions by encoding a source cell state (𝑠_0_) and decoding an experimentally observed downstream target state (*s*_*s*′_), with predictions conditioned on all preceding observed cell states, with attention capturing gene–gene dependencies and their changes across transitions. **b,** Applications of PerturbGen: cell state-conditioned generation of gene expression at specified target states, identification of *de novo* context-specific gene programs from learned gene embeddings, and *in silico* perturbation mapping to organize perturbations by predicted transcriptional responses, with resulting atlases available for interactive exploration (https://cellatlas.io/perturbgen). **c,** Pre-training corpus composition spanning embryonic, fetal and postnatal stages; pie charts indicate the contribution of public and private datasets within each developmental stage (Supplementary Table 1). **d,** Distribution of cells across annotated disease categories in the pre-training corpus.

This framework supports three downstream applications (Fig. 1b). First, PerturbGen predicts gene expression at specified target states, allowing inference of intermediate and future cell states. Second, learned gene embeddings can be aggregated across biological covariates, such as time, lineage, or developmental stage, to identify *de novo*, context-specific gene programs beyond predefined pathway annotations. Third, PerturbGen enables *in silico* perturbation analysis by simulating genetic interventions across cellular states. Scaling these simulations across genes yields perturbation atlases in which perturbations with similar transcriptional effects cluster together. We define these clusters as PIPs, which facilitate systematic identification of established regulators and discovery of previously unrecognized drivers of cell state transitions.

To enable generalization across diverse biological systems and support cell state-aware predictions, we pre-trained the PerturbGen encoder on approximately 107 million single-cell transcriptomes spanning over 100 tissues, diverse diseases and experimental conditions (Fig. 1c,d; Supplementary Table 1). This large-scale pre-training allows the model to learn shared transcriptional programs and gene–gene dependencies that recur across biological contexts. The pre-training corpus included a broad range of postnatal and adult tissues, alongside embryonic and fetal datasets that exhibit high cellular plasticity and densely sampled developmental transitions that are underrepresented in publicly available resources. This selection exposed the model to a broader spectrum of state changes than are typically observed in mature tissues. The pre-trained model was subsequently adapted to stage-resolved data to capture context-specific, state-dependent gene expression dynamics (see Methods).

To benchmark PerturbGen’s generative capability, we evaluated its ability to predict temporal gene expression changes across three time-resolved scRNA-seq datasets spanning immune activation and embryonic development. We compared PerturbGen with representative generative dynamics models, including neural ODE^37^ and flow-matching approaches^41,49^. Across datasets, PerturbGen achieved the best performance for unseen time point prediction and faithfully recovered stage-specific gene expression signatures (Supplementary Fig. 1a,b) Ablation analyses further examined the impact of generative decoding and model parameter choices on performance with tokenized single-cell data (Supplementary Fig. 2a-h), demonstrating that explicitly conditioning on source and contextual states improves generative fidelity and enables accurate modeling of cellular trajectories across time.

### PerturbGen predicts how cytokine perturbation reshapes downstream immune programs during human *in vivo* LPS challenge

Lipopolysaccharide (LPS) is a well-established model of innate immune activation relevant to sepsis-related pathology^50–52^, and induces a rapid systemic inflammatory response, particularly in monocytes, macrophages and B cells. To capture these dynamics *in vivo*, we generated longitudinal scRNA-seq profiles of peripheral blood mononuclear cells (PBMCs) from healthy volunteers undergoing intravenous LPS challenge (six donors; 23 blood samples, Methods). Samples were collected at baseline and at 90 min, 6 h and 10 h following LPS administration, with pre-infusion samples serving as matched controls (Fig. 2a). Across all samples, we profiled a total of 119,360 cells, classified into 30 immune cell types using CellTypist^53^ (Supplementary Fig. 3a-c). With this study, we release the complete dataset, combining newly generated data with a publicly available single-cell PBMC dataset from an in vivo LPS challenge^54^, Cell-type annotations were harmonized across the two datasets, yielding a combined longitudinal atlas of 223,478 cells. This longitudinal design enables interrogation of transcriptional programs underlying dynamic immune state transitions in their physiological context and evaluation of PerturbGen’s ability to model these transitions and perform *in silico* perturbations.

**Fig 2.**
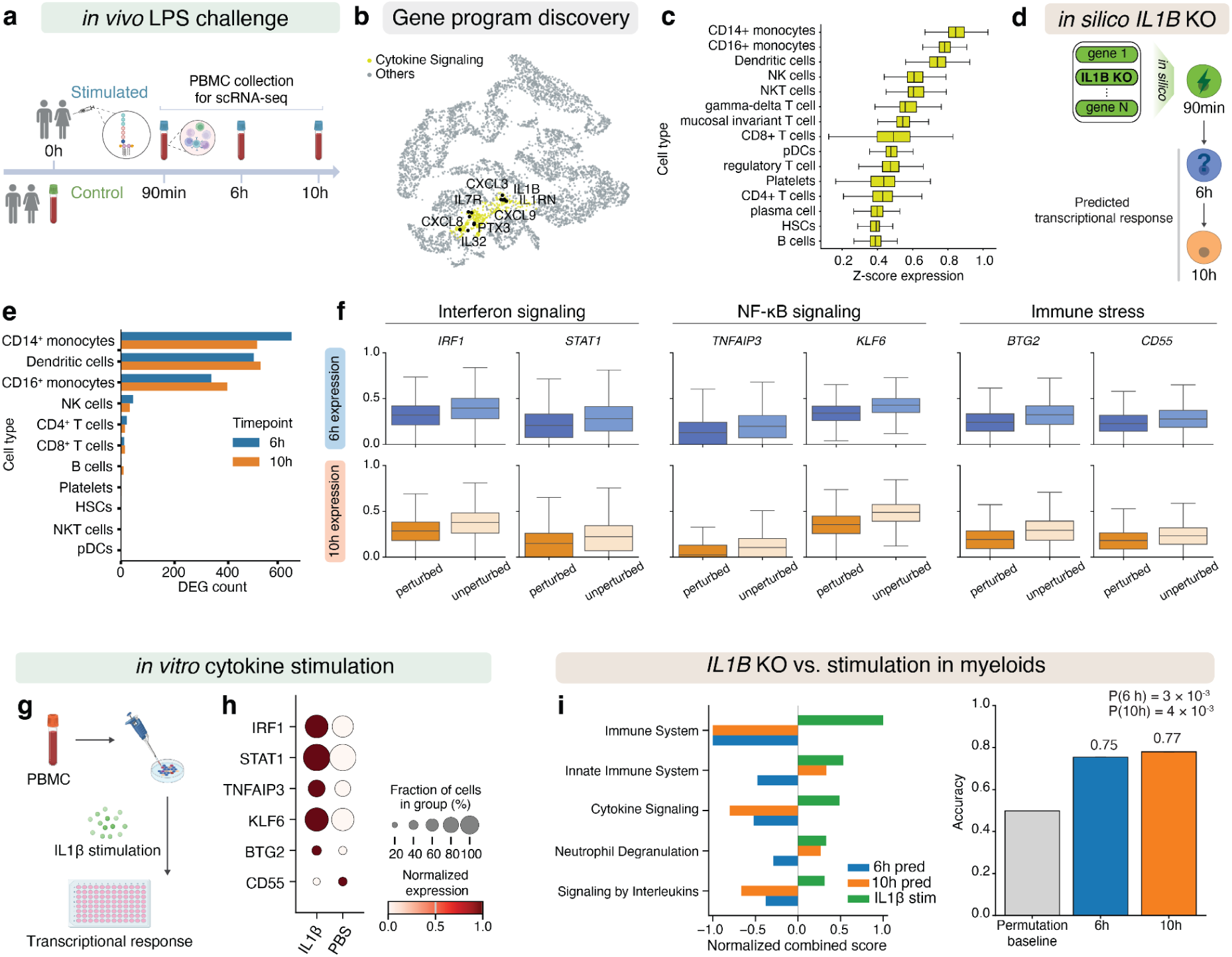
PerturbGen predicts the downstream immune responses to loss of IL1B-mediated signaling during an *in vivo* LPS challenge. **a,** Experimental design for an *in vivo* LPS challenge in healthy participants. Peripheral blood mononuclear cells (PBMCs; n=119,360) were profiled by scRNA-seq at 90 min, 6 h, and 10 h post-injection, with matched controls. **b,** UMAP visualization of aggregated gene embeddings across time points. Each point represents a gene, with a cytokine-associated cluster and selected genes highlighted. **c,** Box plots of the z-scored cytokine/chemokine program score across immune cell types. **d,** Schematic of the *in silico* perturbation experiment: *IL1B* expression was computationally knocked out in cells at 90 min and transcriptional states predicted at 6 h and 10 h. **e,** Cell type-specific perturbation effect quantified as the number of significantly differentially expressed genes (DEGs) between perturbed and unperturbed cells at 6 h (top) and 10 h (bottom). **f,** Representative inflammatory and innate immune regulators with lower predicted expression in perturbed compared with unperturbed cells at 6 h and 10 h. Box plots show predicted min-max normalized expression in unperturbed and perturbed cells at 6 h (top) and 10 h (bottom). **g,** External validation using an independent large-scale arrayed cytokine stimulation screen. PBMCs were stimulated with IL1β or PBS and transcriptionally profiled. **h,** Dot plots show normalized expression of representative inflammatory regulators in myeloid cells, with dot size indicating the fraction of expressing cells. **i,** Mirror plot of pathway enrichment in myeloid lineages. Pathways downregulated following *in silico IL1B* knockout (6 h and 10 h predictions) are compared to pathways upregulated following IL1β stimulation in the external dataset. **j,** Pathway-level reversal accuracy, measuring concordance in the direction of pathway regulation between *IL1B* knockout prediction and IL1β responses, shown for 6 h predictions, 10 h predictions and a random permutation baseline (empirical *P* value). Box plots indicate median (center line), interquartile range (box) and range (whiskers).

We applied PerturbGen to the longitudinal LPS atlas to model the endogenous immune response and examine how gene perturbations alter downstream transcriptional responses across immune cell types. We first captured transcriptional dynamics across the *in vivo* LPS response by deriving representations for each gene. Clustering these representations across time points revealed distinct gene programs of immune activation with shared temporal patterns (Fig. 2b; Supplementary Fig. 4a,b). Early inflammatory responses are orchestrated by cytokine signaling networks, prominently including IL-1β-mediated pathways^55^. Consistent with this, clustering of gene representations identified a cytokine-associated program enriched for genes involved in *IL-1β* signaling, including *IL1B*, *IL1RN* and *CXCL8*, indicating that PerturbGen captured known early inflammatory pathways. Gene expression analysis revealed that this IL-1β-mediated program was preferentially enriched in CD14⁺ monocytes, CD16⁺ monocytes and dendritic cells (DCs) (Fig. 2c). Within these myeloid subsets, IL-1β-mediated program activity increased rapidly following LPS exposure, with the highest scores observed at 90 min and 6 h (Supplementary Fig. 5a). *IL1B* expression itself peaked at 90 min and declined at later time points (Supplementary Fig. 5b), suggesting a temporal window during which cytokine signaling influences downstream cellular states. Consistent with its role as an early NF-κB–dependent TLR4 target^56^, IL-1β initiates acute inflammatory programs and shapes downstream myeloid states, making it a biologically grounded proof-of-principle perturbation.

Given the sharp, transient peak of *IL1B* at 90 min, we leveraged PerturbGen to perform an *in silico* knockout in LPS-stimulated cells at this early time point and predict transcriptomic states at 6 h and 10 h (Fig. 2d, Supplementary Table 12), quantifying the effects of the perturbation as the number of differentially expressed genes (DEGs) between perturbed and unperturbed cells. Concordant with the enrichment of the cytokine-associated gene program in CD14⁺ or CD16⁺ monocytes and dendritic cells, and consistent with the established role of IL-1β in myeloid cells^57^, the largest number of DEGs after perturbation was observed in myeloid populations (Fig. 2e). As perturbation-induced transcriptional changes were concentrated within myeloid subsets (monocytes and DCs), subsequent analyses focused on these subsets. Pathway enrichment analyses of DEGs highlighted decreased enrichment of cytokine signaling, innate immune response pathways, and interferon-associated programs following *in silico IL1B* KO (Fig. 2f; Supplementary Fig. 6a,b). This included genes linked to interferon-associated responses (for example *IRF1*, *STAT1*)^58^, NF-κB pathway components (for example *KFL6*, *TNFAIP3*)^59,60^, and AP-1-associated transcriptional regulator (for example *JUNB, FOS*)^61,62^. Reduced predicted expression was also observed for genes involved in immune cell trafficking and activation (for example *CXCR4*, *RHOB*)^63,64^ as well as immune-related stress genes (*BTG2*, *CD55*)^65,66^ (Fig. 2f; Supplementary Fig. 7a).

To assess the biological plausibility of our perturbation, we compared our predicted transcriptomic changes following *in silico IL1B* perturbation with an independent dataset in which primary human PBMCs were stimulated with IL-1β^67^, reflecting the opposite directionality of gene perturbation (Fig. 2g). At the gene level, multiple regulators of the innate inflammatory response were predicted to decrease following *IL1B* perturbation. Conversely, many of the same genes showed increased expression following IL-1β stimulation in the external dataset (Fig. 2f,h). At the pathway-level, pathways upregulated following IL-1β stimulation showed decreased enrichment in *IL1B* perturbation predictions at 6 h and 10 h (Fig. 2i). Similar pathway-level changes were observed following *in silico* perturbation of the IL-1 receptor (Supplementary Fig. 8a, b). Pathway-level accuracy of this reversal reached 75% at 6 h and 77% at 10 h, compared with a random permutation baseline generated by reassigning predicted gene-level effect directions while preserving pathway structure (Methods; empirical *P* = 0.003 for 6 h and empirical *P =* 0.004 for 10 h; Fig. 2j; Supplementary Fig. 9a, b).

Overall, these findings provide a proof-of-principle that perturbing an early-response cytokine during a human immune challenge can predict sustained downstream shifts in immune cell state. *In silico IL1B* perturbation attenuated inflammatory programmes, opposing the effects of IL-1β stimulation observed in an independent dataset. Together, this supports that PerturbGen recapitulates established immune biology and generalizes across experimental contexts.

### A lineage-aware *in silico* perturbation atlas recovers canonical cell states and predicts disease phenotypes in human hematopoiesis

To evaluate PerturbGen in a dynamic developmental system beyond acute inflammation, we profiled human hematopoiesis at single-cell resolution. Hematopoietic stem and progenitor cells (HSPCs) comprise lifelong, self-renewing stem cells and lineage-committed progenitors that generate diverse blood cell states^68^. We generated an atlas of human CD34^+^ HSPCs (n = 98,266) spanning yolk sac (6–7 post-conception week(PCW)), fetal liver (6–17 PCW), fetal bone marrow (14–17 PCW) and cord blood through pediatric (1–16 years) and adult bone marrow (29–91 years) (Fig. 3a), and used it to train PerturbGen to capture the transcriptomic networks underlying age- and tissue-specific hematopoietic decision making. Using CITE-seq^69^, we identified 23 HSPC populations ranging from phenotypic long-term HSCs to lineage-committed progenitors across megakaryocyte (MK), erythroid, eosinophil/basophil/mast cell, B-lymphoid, natural killer/T, monocyte/macrophage, dendritic and plasmacytoid dendritic lineages (Fig. 3b; Supplementary Fig. 10a,b and 11; Methods). We observed the expected enrichment for macrophage production in yolk sac^70^, erythropoiesis in fetal liver^71–73^ and B-lymphopoiesis in fetal and pediatric bone marrow^4,72,74^ (Supplementary Fig. 10c,d). This dataset, therefore, provides a comprehensive platform to study age- and lineage-specific hematopoietic regulation across the human lifespan.

**Fig 3.**
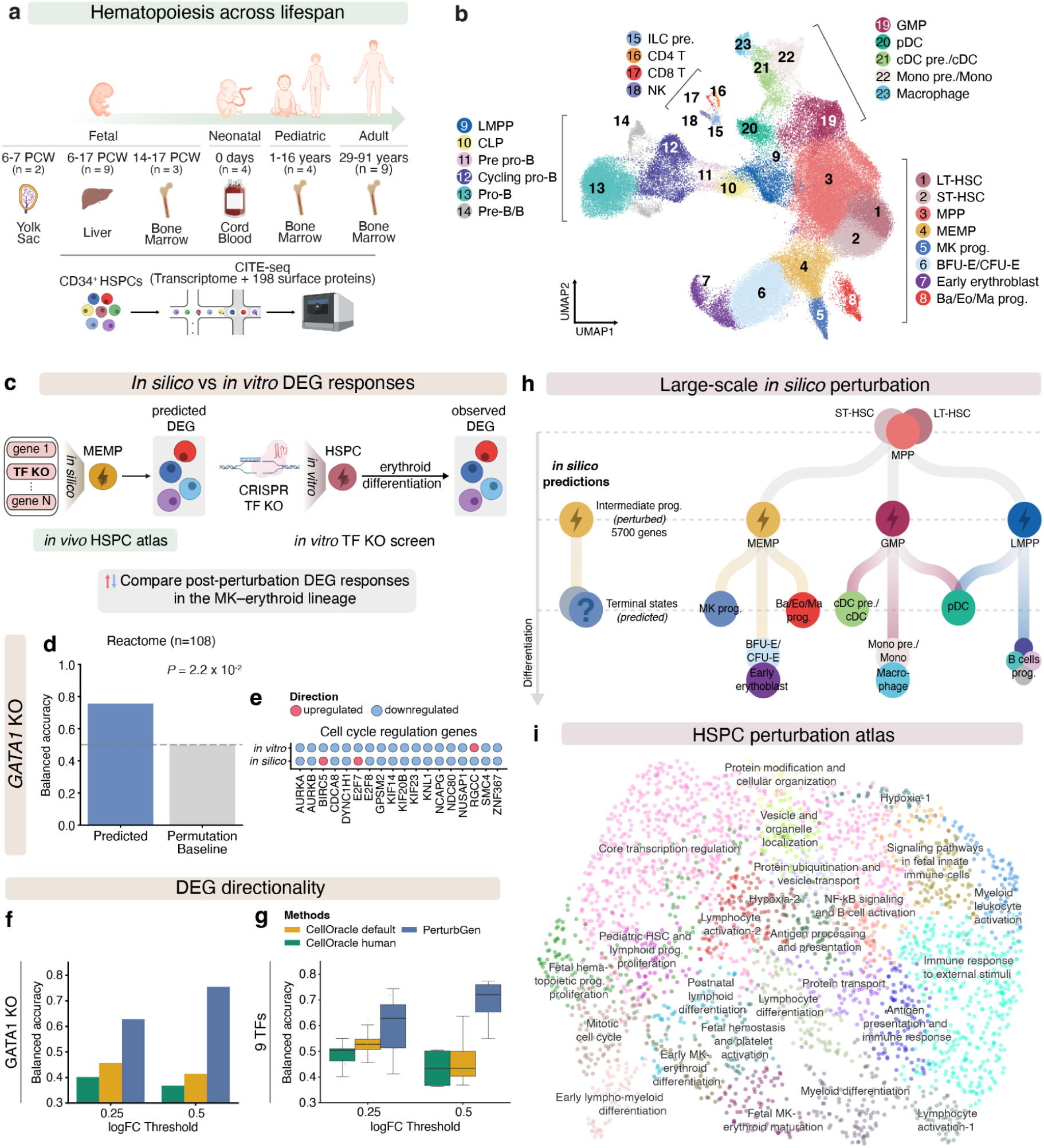
Systematic *in silico* perturbation simulation during human hematopoiesis. **a,** Study design schematic depicting CITE-seq profiling of CD34^+^ human hematopoietic stem and progenitor cells (HSPCs) spanning fetal to adult tissues (n = 31 samples from 26 donors). **b,** Joint transcriptomic and proteomic profiling of 98,266 cells across myeloid, lymphoid and megakaryocyte–erythroid–mast cell lineages. Abbreviations for hematopoietic populations follow standard nomenclature (e.g., HSC, MPP, LMPP, GMP, MEMP; see Methods). **c,** Evaluation strategy comparing *in silico* gene perturbations simulated within the *in vivo* HSPC atlas with *in vitro* perturbations measured using pooled CRISPR transcription factor (TF) knockouts (KO) in lineage-committed progenitor states across ten TFs. **d,** Pathway-level concordance between *in silico* and *in vitro GATA1* KO, quantified using normalized enrichment scores (NES) from gene set enrichment analysis (GSEA) of Reactome. Model performance was evaluated using balanced accuracy for pathway directionality (up- versus down-regulated) and compared to a permutation-based null distribution (empirical *P* value shown). **e,** Concordance in directionality of GSEA lead genes involved in positive regulation of the cell cycle. Dots are colored by log₂ fold-change sign (red, positive; blue, negative). **f,** Prediction performance for *GATA1* KO, showing directionality balanced accuracy across different log₂ fold-change thresholds of 0.25 or 0.5 (adjusted *P* value < 0.05), comparing PerturbGen with CellOracle using default or human ATAC-derived gene regulatory networks. **g,** Summary of benchmarking performance across nine TFs, evaluated across log₂ fold-change thresholds 0.25 and 0.5. **h,** PerturbGen training strategy in which cells are paired according to the hematopoietic lineage hierarchy, with *in silico* gene KOs applied to intermediate progenitor states, and perturbation effects evaluated in the downstream terminal states. **i,** UMAP visualization of perturbation embeddings in terminal states for 5,700 simulated gene perturbations. Each point represents a single gene perturbation, mean-pooled across cells with detectable expression of the perturbed gene. PIPs are annotated based on shared functional and lineage-specific effects (Methods; Supplementary Fig. 14a-f).

To establish the validity of PerturbGen genetic perturbation predictions for human hematopoiesis, we evaluated its performance against a multi-omic CRISPR transcription factor (TF) KO dataset in human HSPCs^75^. In this reference dataset, 19 TFs, established regulators of hematopoietic stem and progenitor cell function, were perturbed in HSPCs before culturing in erythroid-promoting culture conditions. Of the 19 TFs, nine exhibited robust transcriptional shifts following experimental perturbation (Methods; Supplementary Fig. 12b), as determined by minimum DEG threshold across log₂ fold-change cutoffs. We then performed *in silico* perturbations of the same nine genes within our atlas of human hematopoiesis. To ensure comparability with this CRISPR dataset generated under an erythroid-promoting culture condition, we focused on megakaryocyte–erythroid–mast cell progenitors (MEMPs), and evaluated the predicted transcriptional effects on erythroid lineage commitment in the downstream populations (Fig. 3c), following cross-dataset cell type alignment using a similarity metric (Supplementary Fig. 12a). We first compared pathway-level responses following experimental and *in silico* KO of *GATA1,* a canonical erythroid-megakaryocyte master regulator^76^. PerturbGen correctly predicted the direction of regulation of 76% of Reactome gene sets (n = 108). This agreement was significantly greater than expected under a randomized baseline in which gene rankings were permuted prior to pathway analysis (empirical *P* = 0.022; Fig. 3d, Supplementary Fig. 13a, b). Consistent with these pathway-level results, genes involved in positive regulation of the cell cycle were downregulated in both settings, with concordant directionality observed for 15 of 18 genes (Fig. 3e).

We next benchmarked the generative performance of PerturbGen against CellOracle^28^, a GRN-based framework that predicts perturbation effects using ATAC-derived regulatory networks. Unlike CellOracle, which models perturbation effects within individual cell states, PerturbGen simulates perturbations in precursor populations and predicts their downstream effects along differentiation trajectories. Across log₂fold-change thresholds of 0.25 or 0.5 (adjusted *P* < 0.05), PerturbGen consistently outperformed CellOracle in predicting DEG directionality. As *GATA1* exhibited the strongest transcriptional response upon experimental perturbation, we first evaluated model performance for *GATA1* KO. PerturbGen outperformed CellOracle for *GATA1* in both balanced accuracy and F1 macro (Fig. 3f, Supplementary Fig. 13c). This trend was consistent across all nine transcription factor KOs, with PerturbGen achieving a mean balanced accuracy of 62.8-72.0% and F1 macro scores of 53.4-55.6%, compared to 43.5–52.7% accuracy and 22.9–28.9% F1 macro for CellOracle (Fig. 3g, Supplementary Fig. 13d). These results highlight PerturbGen’s ability to accurately model downstream transcriptional propagation of genetic perturbations beyond individual cell states.

In light of these results, we next upscaled to identify genes and gene programs associated with age- and lineage-specific cell state transitions in human hematopoiesis in an unbiased manner. We thus performed an *in silico* perturbation screen, targeting oligopotent intermediate progenitor states (lymphoid-primed multipotent progenitors (LMPPs), granulocyte–macrophage progenitors (GMPs) and megakaryocyte–erythroid–mast cell progenitors (MEMPs)) and inferred perturbation effects on downstream terminal, lineage-committed progenitor states (Fig. 3h). In total, 5,700 genes were evaluated, including highly variable genes and transcribed transcription factors; of which, 3,108 passed quality control and were embedded into a *perturbation map* (Methods). Unsupervised Leiden clustering partitioned this map into distinct PIPs, each capturing shared transcriptional responses to genetic perturbation (Fig. 3i). These programs exhibited distinct age- and/or lineage-specific expression patterns and distinct enrichments for functional gene sets. For instance, one PIP showed high activity in postnatal HSCs and early lymphoid progenitors (Supplementary Fig. 14a). This PIP included established regulators of lymphoid differentiation, such as *FLT3*^77^, and showed enrichment for gene sets linked to lymphoid development (Supplementary Fig. 14b). We therefore annotated this program as “postnatal lymphoid differentiation”. Interestingly, we identified two distinct cell cycle programs with age-and lineage-specific expression patterns, annotated as “pediatric HSC and lymphoid progenitor proliferation” and “fetal hematopoietic progenitor proliferation”. While both PIPs showed an enrichment of cell cycle-related gene sets, the former exhibited high activity in pediatric HSCs and lymphoid progenitors (Supplementary Fig. 14c,d), whereas the latter was highly expressed in fetal HSPCs (Supplementary Fig. 14e,f), suggesting the presence of distinct cell cycle mechanisms that operate differently across developmental stages of human hematopoiesis. Together, these results demonstrate that systematic *in silico* perturbation modeling identifies context-specific functional regulatory programs based on shared downstream transcriptional cascades.

Having established a large-scale hematopoiesis perturbation atlas, we next asked whether PIPs identified by PerturbGen were associated with human hematological traits and disorders. Because genes within shared functional modules frequently underlie overlapping diseases or phenotypes^78^, we linked PIPs to human genetic and disease associations using the Open Targets (OT) Platform, which integrates genetic and functional evidence for target–disease associations^79^. PIPs were mapped to blood-related traits^80^ using Locus-to-Gene (L2G) scores from fine-mapped genome-wide association study (GWAS) loci, and rare-disease associations curated from OT supported by ClinVar^81^ variant annotations and PanelApp^82^ expert curated gene-disease evidence (Fig. 4a). Lineage-specific PIPs showed enriched phenotypic associations with corresponding hematopoietic lineages. Genes within the myeloid differentiation PIP showed multiple associations with monocyte and granulocyte GWAS traits (43 genes; Fig. 4b). Likewise, multiple genes within the lymphocyte activation PIPs were associated with the lymphocyte count and related lymphocyte traits (34 genes; Supplementary Fig. 15b). Furthermore, the early megakaryocyte-erythroid differentiation PIPs showed associations with platelet and red blood cell traits (43 genes; Supplementary Fig. 15a). Beyond quantitative traits, lineage-specific PIPs exhibited associations with relevant hematological disorders. The early megakaryocyte–erythroid PIP was enriched for MK/platelet disorder genes (9 observed vs 1.3 expected; OR=8.9; global FDR=3.7×10⁻⁴; Supplementary Fig. 15b), while the lymphocyte activation PIP was enriched for immunodeficiency genes (11 observed vs 2.0 expected; OR=7.1; global FDR=3.7×10⁻⁴; Supplementary Fig. 15d). Importantly, while these PIPs included well-established regulators of hematopoietic lineage differentiation and blood/immune disorders, such as *TAL1* in erythropoiesis^83^ and *EBF1* for B-lymphoid development^84^, they also comprised numerous previously unrecognized genes that were nonetheless associated with lineage-specific hematological traits and disorders (Fig. 4b, Supplementary Fig. 15a,b). Together, these results demonstrate that PerturbGen identifies gene programs that are both biologically coherent and translationally meaningful.

**Fig 4.**
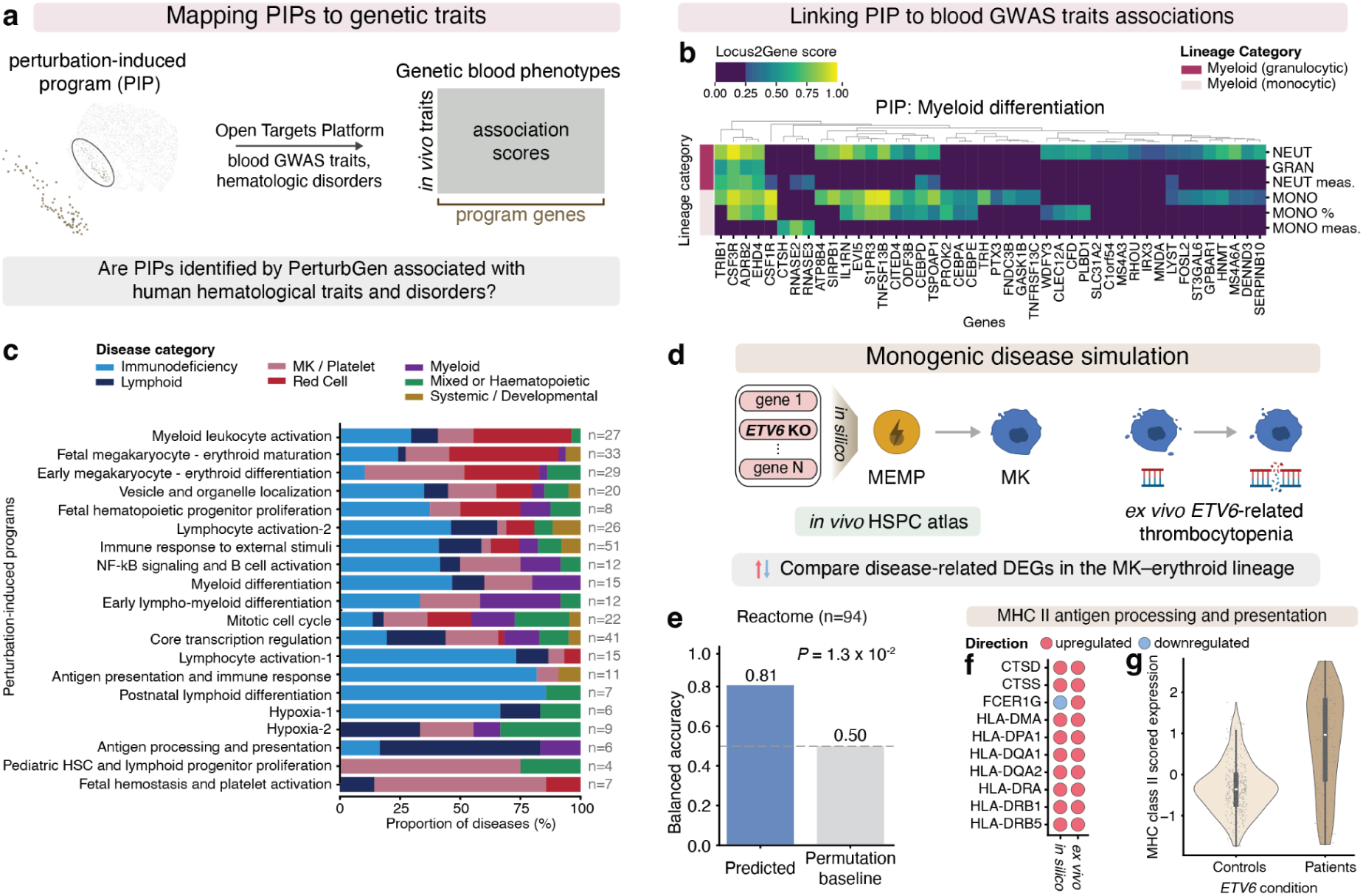
Linking perturbation-induced programs to blood traits and monogenic disease. **a,** Schematic illustrating the mapping of PIPs to *in vivo* blood-related traits and diseases using the Open Targets platform, integrating genome-wide association study (GWAS)-derived Locus-to-Gene (L2G) scores and rare-disease gene annotations from ClinVar and PanelApp. **b,** Clustered heatmaps showing associations between genes and hematological GWAS traits for a representative PIP corresponding to myeloid differentiation. Associations are colored by the Open Targets L2G scores (≥0.25), and row annotations indicate lineage category for each trait. **c,** Proportion of genetically supported diseases associated with each PIP, stratified by disease category. The total number of diseases is indicated at the end of each row (n=x). **d,** Schematic of monogenic disease simulation. An *in silico ETV6* knockout in megakaryocyte progenitors (MK prog.) within the *in vivo* hematopoietic stem and progenitor cell landscape is compared with *ex vivo*-derived MK prog. from patients with *ETV6*-associated thrombocytopenia. **e,** Comparison of pathway-level gene set enrichment analysis (GSEA) between *in silico* perturbation prediction and patient-derived *ETV6* thrombocytopenia data. Model performance is assessed using balanced accuracy and contrasted with a permutation-based baseline; *P* indicates the empirical *P* value. **f,** Dot plots showing concordant directionality of lead genes from GSEA for antigen processing and presentation pathway. Dot color indicates log₂ fold-change direction (red, upregulated; blue, downregulated). **g,** Distribution of MHC class II-related gene module scores in a lineage-restricted megakaryocyte population, comparing controls with patients with ETV6-associated thrombocytopenia (control n=2, patient n=2).

To further assess the translational relevance of our hematopoiesis perturbation atlas, we explored whether PerturbGen can recapitulate transcriptomic alterations observed in monogenic blood disorders. Owing to the limited availability of single-cell datasets for rare monogenic blood disorders, we focused on a study of *ETV6*-related thrombocytopenia^87^, in which *ex vivo*-expanded CD34^+^ HSPCs from two patients were differentiated into megakaryocytes and subjected to scRNA-seq. We compared the predicted perturbation effects in *in silico ETV6* KO in megakaryocyte progenitors within our hematopoiesis perturbation atlas with disease-associated transcriptional alterations observed in patient megakaryocyte progenitors (Fig. 4d). At the pathway level, this comparison revealed a highly shared, non-random signal (empirical *P* = 0.00128), with 81% directional concordance of gene sets enriched for disease relevance (Fig. 4e; Supplementary Fig. 16a). Concordant pathways included upregulation of antigen processing and presentation, including MHC class II-related pathways, alongside downregulation of platelet-associated processes. In contrast, cell cycle-related gene sets were predominantly downregulated in patient samples but showed mixed responses in the model predictions (Supplementary Fig. 16b, c), which may reflect differences in baseline cell cycle activity between healthy reference cells and *ex vivo*-expanded patient cells (Supplementary Fig. 16d). At the gene level, PerturbGen recapitulated key disease-associated transcriptional changes, including upregulation of antigen processing and presentation pathways (9/10 concordant genes; Fig. 4f), as well as downregulation of platelet-related genes (6/9 concordant genes) and cell-cell junction organization (5/5 concordant genes; Supplementary Fig. 16e, f). To exclude the possibility that increased expression of MHC class II-related genes reflected a shift in cell composition towards myeloid lineages, we further restricted our analysis to cells expressing megakaryocyte (MK) markers while lacking expression of myeloid-associated genes. Consistent with the full analysis, this filtered population showed an increase in MHC class II-related genes and decreased expression of MK and platelet-associated genes (Fig. 4g; Supplementary Fig. 17a-d). These effects were consistent across both patients and were recapitulated in pseudobulk analyses, indicating that they were not driven by compositional differences (Supplementary Fig. 17e, f). Collectively, these findings illustrate the ability of PerturbGen to recapitulate disease-associated transcriptional signatures observed in monogenic disorders.

Together, these analyses leverage PerturbGen to generate a trajectory-aware *in silico* perturbation atlas of human hematopoiesis, systematically linking genetic perturbations in progenitor states to downstream lineage outcomes. Our predictions could be organized into age- and lineage-specific PIPs, which were anchored to human genetic traits and model a human monogenic blood disorder.

### PerturbGen prioritizes Wnt activation to enhance stromal maturation in skin organoids towards *in vivo* fibroblast states

We then extended our use of PerturbGen to prioritize molecular interventions in a complex multicellular organoid system. A major challenge in developing organoids is to drive these *in vitro* systems to more closely recapitulate human biology, and to identify specific biological, chemical or physical cues that induce cell states observed *in vivo*^88,89^. We therefore applied PerturbGen to human induced pluripotent stem cell (iPSC)-derived skin organoids. Although iPSC-derived skin organoids faithfully recapitulate major aspects of epidermal and dermal development, as well as spatial organization of human skin^88,89,6,90^, some organoid cell types do not reach primary human skin-like states. For example, human adult skin contains distinct fibroblast subsets that have been identified in microanatomical niches within the human skin, with putative roles in hair follicle differentiation and immunoregulation^91^. However, despite containing a large population of stromal cells^6,90^, human skin organoids generated from iPSCs are typically cystic and lack several *in vivo* microenvironmental cues that may support stromal maturation, such as stable tissue anchoring, sparse vasculature and no immune cells. As such, they lack the full spectrum of dermal fibroblast heterogeneity observed *in vivo*.

We therefore applied PerturbGen to identify cues for promoting fibroblast maturation in human skin organoids. Taking our previously published human skin organoid scRNA-seq dataset^6,90^ (Fig. 5a), we performed an *in silico* perturbation screen across 5,050 genes, comprising highly variable genes together with additional genes implicated in developmental processes of the organoid. Our *in silico* perturbations were targeted on day 6 of organoid differentiation as the highest number of cells in the early stages of differentiation, aiming to identify putative molecular targets at the earliest profiled time point to alter cell states at later time points (Supplementary Fig. 18a). For each day of skin organoid differentiation, predicted transcriptional responses after our *in silico* screen were embedded into day-specific perturbation maps (Supplementary Fig. 18b). We used unsupervised clustering to identify PIPs at day 29, reflecting an early time point in skin organoid development^6,90^ and determining which molecular perturbations at day 6 which would promote stromal differentiation and dermal fibroblast maturation *in vitro*. We annotated six PIPs, including: interferon response, protein synthesis, cell-cycle regulation, extracellular matrix remodeling, and additional developmental pathways including proliferation and keratinization (Fig. 5b; Supplementary Table 19).

**Fig 5.**
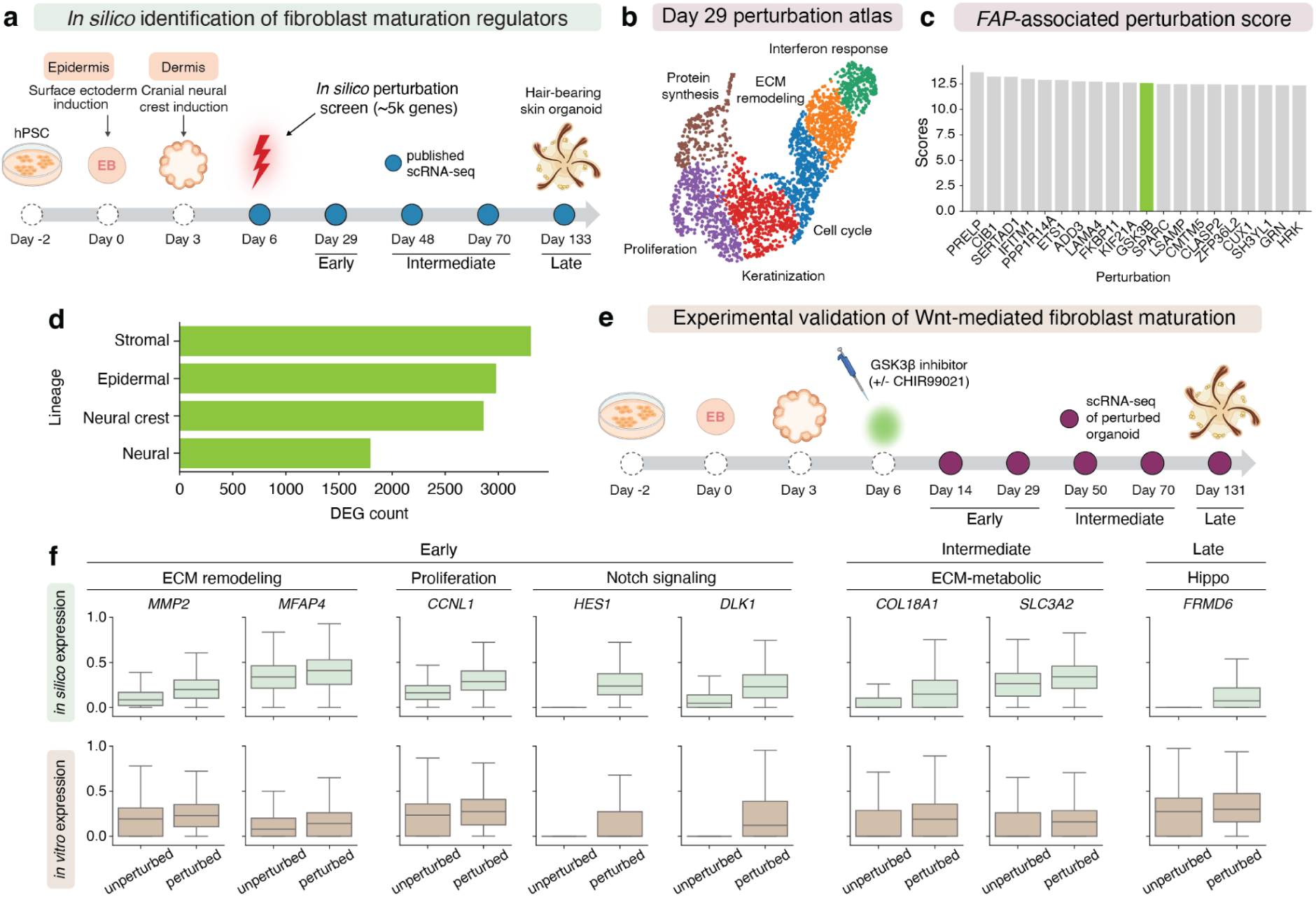
PerturbGen-guided identification of regulators of stromal maturation in skin organoids. **a,** Schematic overview of the published skin organoid developmental dataset used for PerturbGen analysis. An *in silico* perturbation screen was conducted spanning ∼5,000 genes at an early time point, and transcriptional states were predicted across early (D0–45), mid (D45–90), and late (D90–135) of organoid development. **b,** UMAP visualization of the perturbation atlas at day 29, in which each point represents a single gene perturbation aggregated across all cell types colored by annotated perturbation-induced programs. **c,** Bar plot visualization showing the top-scoring perturbations ranked by predicted *FAP* expression change within the stromal lineage at day 29. **d,** Lineage-specific perturbation effects following *GSK3B* perturbation, quantified as the number of DEGs aggregated across developmental time points. **e,** Schematic of experimental *in vitro* Wnt activation used for validation. Skin organoids were treated with a GSK3β inhibitor (CHIR99021) at day 6 of differentiation, and profiled by scRNA-seq at early, mid, and late stages. **f,** Box plots showing min-max normalized expression changes of genes associated with stromal maturation across developmental stages. The top row shows PerturbGen predictions, and the bottom row shows experimentally measured expression following *in vitro* Wnt activation. Box plots indicate median (center line), interquartile range (box) and range (whiskers).

We then prioritised *in silico* perturbations at day 6 of human skin organoid differentiation that were predicted to enhance fibroblast maturation at day 29. We used predicted expression of fibroblast activation protein (*FAP*) as a quantitative readout, due to recent evidence indicating its expression as a pan-tissue fibroblast marker and its upregulation in activated dermal fibroblast states^91,92^. We ranked all day 6 genetic perturbations by their predicted effect on altering *FAP* expression in stromal populations at day 29 (Fig. 5c). The top 20 perturbations showed comparable predicted effects on *FAP* expression, and contained known regulators of fibroblast differentiation, activation or polarisation including *PRELP*^93,94^*, ETS*^95^ and *LAMA4*^96^. Amongst the top candidates was the negative regulator of canonical Wnt/β-catenin signaling, *GSK3B*^97^. Importantly, a selective small molecule inhibitor of GSK3β, CHIR99021, has been shown to promote human skin organoid growth^98^. Further analysis of *in silico GSK3B* perturbation by quantifying DEGs across all major cell lineages revealed the largest number of DEGs to be within the stromal lineage (Fig. 5d; Supplementary Fig. 19a), further prioritising this molecule as a candidate for biological validation.

To experimentally test PerturbGen’s prediction that modulation of GSK3β would promote stromal differentiation in human skin organoids, we generated new scRNA-seq data after culturing organoids *(n =* 3-6 organoids per group) at day 6 with and without CHIR99021 (Fig. 5e and Supplementary Fig. 20a-c). To compare these biological validation experiments to predictions from PerturbGen *in silico* perturbations, we grouped time points into three stages of the organoid culture, broadly corresponding to major morphological and molecular milestones reported in human skin organoid systems, including early (D0-45, lineage induction and tissue organisation), intermediate (D45-90, early hair follicular morphogenesis), and late (D90-135, increasing appendage complexity and follicular maturation) stages^88,89,6,90^. Predicted transcriptional changes in stromal populations following *in silico GSK3B* perturbation were consistent with those observed in CHIR99021-treated skin organoids. In both *in silico* data and *in vitro* validation, upregulated genes included in ECM remodelling (*MMP2*, *MFAP4*), proliferation (*CCNL1*) and Notch signaling (*HES1*, *DLK1*) at the early stage of skin organoid differentiation, basement membrane deposition (*COL18A1*) and amino acid transport (*SLC3A2*) at the intermediate stage, and the Hippo pathway signaling regulator *FRMD6* at the late stage (Fig. 5f). The concordance in DEGs between *in silico* perturbations and *in vitro* data suggests that PerturbGen captures aspects of the observed response and motivated further profiling of how GSK3β inhibition influences skin organoid stromal differentiation.

### GSK3β inhibition biases skin organoid stroma toward a primary fetal skin-like state

Given the stromal responses to *in silico* GSK3B inhibition in human skin organoids, we further interrogated our *in vitro* GSK3β inhibition scRNA-seq data, again grouping the data into early, intermediate or late stages (Fig. 6a). A total of 104,554 cells were pooled and profiled from unperturbed and GSK3β-inhibited conditions, resolving into six major lineages represented in prior scRNA-seq studies of human skin organoids, including epidermal, neural crest, muscle, endothelial, stromal and adipose lineages (Fig. 6b, c). Both CHIR99021 treatment and *in silico GSK3B* perturbation indicated that the most pronounced transcriptional changes occurred in stromal cells at the early time point as quantified by the highest number of DEGs (Supplementary Fig. S21a, Methods). Focused analysis of stromal cells, which exhibited the highest number of DEGs following *in silico* perturbation of *GSK3B*, resolved seven transcriptionally distinct stromal states corresponding to known differentiation stages in hair follicle stromal cell development^88^, beginning in early fibroblasts and progressing to pre-dermal condensate, dermal condensate (DC), and dermal papilla (DP) cell states (Fig. 6d,e). Pseudotime analysis of the CHIR99021-treated stromal cell subset predicted a continuous differentiation trajectory originating from early fibroblasts and progressing toward DC and DP-like states (Fig. 6e), as expected.

**Fig 6.**
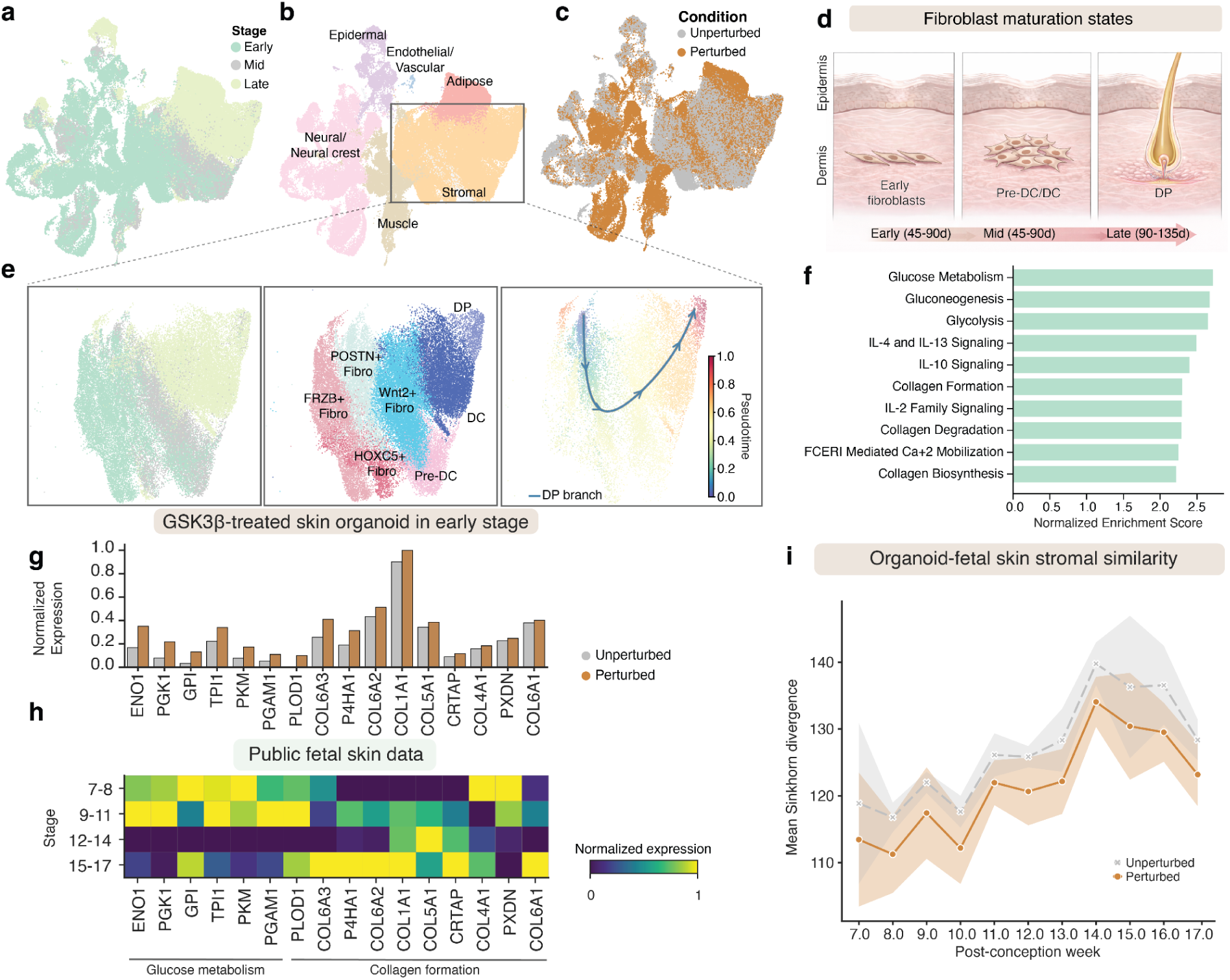
GSK3β inhibition promotes fetal-like stromal features in skin organoids. **a-c,** UMAP visualization of single-cell transcriptome from skin organoids exposed to GSK3β inhibition, colored by **a)** developmental time points (early, mid, and late), **b)** broad cell lineages labels and **c)** experimental conditions. **d,** Schematic of stromal maturation during skin development, illustrating progression from early fibroblasts through a pre-dermal condensate (pre-DC) state toward dermal papilla (DP) cells. **e,** Subsetted UMAP visualization of stromal cells across developmental stages (left), annotated stromal cell states (middle) and the inferred GSK3β-inhibited developmental trajectory colored by pseudotime (right). **f,** Gene set enrichment analysis of GSK3β-inhibited stromal cells relative to unperturbed controls using Reactome gene sets, summarized by normalized enrichment score (NES). **g,** Bar plots of representative genes associated with glucose metabolism, and collagen formation in GSK3β-inhibited and unperturbed skin organoids at early stages. All the genes passed log₂ fold change > 0; adjusted *P* value < 0.05) calculated by Wilcoxon test. **h,** Matrix plots showing normalized expression of the same genes in human fetal skin stromal across grouped stages. **i,** Mean Sinkhorn divergence between organoid stromal and human fetal skin stromal across PCWs. For each PCW and condition, the mean was calculated across organoid stromal stages, and error bands denote the standard deviation of these stage-specific distances.

Among the pathways positively enriched following CHIR99021 treatment in the early stage (Fig. 6f), we observed a strong representation of gene programmes associated with fetal stromal states including glucose metabolism and collagen formation. This prompted us to examine whether CHIR99021-treated skin organoids adopt transcriptional features resembling human fetal skin *in vivo*. We therefore compared positively enriched pathways in stromal cells, focusing on early time points to capture immediate transcriptional responses, with an independent human fetal skin scRNA-seq dataset spanning 7-17 PCWs^6^. To facilitate this comparison, top pathways enriched after CHIR99021 treatment (Fig. 6f) were assessed in the human fetal skin scRNA-seq dataset subsetted to stromal cells. In the organoids, stromal cells showed up-regulation of genes belonging to pathways enriched after CHIR99021 treatment (Fig. 6g; Supplementary Fig. S22a, b), including key glycolytic enzymes (for example, *ENO1* and *PKM*) and collagen genes (for example, *COL1A1* and *COL6A2*). In the fetal stromal cells, glucose metabolism genes peaked during early developmental stages (7-11 PCWs), whereas collagen formation genes became more prominent at later stages (15-17 PCWs) (Fig. 6h).

Finally, we quantitatively assessed transcriptomic similarity between stromal cells in skin organoid and human fetal skin using Sinkhorn divergence, an entropy-regularized optimal transport metric that quantifies the minimal cost of transforming one gene expression distribution into another, thereby providing a global measure of transcriptional similarity between stroma *in vitro* and *in vivo*^99^. We computed Sinkhorn divergence between stromal cells from CHIR99021-treated or untreated skin organoids, using scRNA-seq data of human fetal skin stroma spanning 7-17 PCWs as a reference (Methods); Sinkhorn divergence is an optimal transport–based distance that summarizes how similar two distributions are, with lower values indicating greater overall transcriptomic similarity. CHIR99021-treated skin organoid stromal cells exhibited reduced divergence relative to fetal skin stroma compared with unperturbed controls (Fig. 6i and Supplementary Fig. S23a, mean Cohen’s d ≈ 1.16), indicating that CHIR99021 treatment shifted the organoid stromal cells towards stromal cell states observed in human skin development *in vivo*.

Using PerturbGen to construct an *in silico* gene perturbation atlas, we identified Wnt activation via GSK3β inhibition as a potential approach to promote stromal differentiation in human skin organoids. Consistent with this prediction, pharmacological inhibition of GSK3β with CHIR99021 induced the expected stromal transcriptional programs and shifted organoid stroma toward fetal skin-like states. Together, these findings suggest that trajectory-aware *in silico* perturbation analyses can help identify experimental strategies to guide complex in vitro tissues toward desired *in vivo* like cell states.

## Discussion

In this study, we present PerturbGen, a generative foundation model for predicting genetic perturbation responses across cellular state transitions. PerturbGen is pre-trained on over 100 million single-cell transcriptomes, and by operationalizing trajectories as experimentally observed discrete states, learns state-to-state transitions with an encoder–decoder transformer trained on tokenised gene expression sequences. This formulation goes beyond within-state perturbation prediction and unifies three capabilities within a single model, whilst maintaining performance in comparison to task-specific approaches. First, it generates trajectory-aware predicted gene expression, conditioned on cell states, to define specified target states, complementing recent generative approaches for temporal and developmental modeling^40,43,44^. Second, it yields interpretable gene embeddings that can be aggregated and clustered across covariates. Thirdly, by conditioning target-state generation on upstream and intermediate states, PerturbGen models how an intervention introduced at a source state propagates to reshape downstream programs and fate-relevant outcomes, enabling systematic construction of *in silico* perturbation atlases without requiring perturbation training data. These three applications turn trajectory-aware prediction into a framework for systematic biological discovery and perturbation prioritisation.

Across newly generated human single-cell datasets spanning innate immunity, hematopoiesis, and skin organoid development, PerturbGen captures that perturbation effects can emerge downstream along trajectories, with consequences that are not limited to the state in which the intervention is introduced. We begin with proof-of-principle experiments in the physiologically relevant human setting of an *in vivo* LPS challenging, a well-established model of endotoxin-driven inflammation and systemic innate immune activation. PerturbGen identified transient *IL1B*-associated program in myeloid cells, successfully predicting known transcriptional responses at later timepoints, including cytokine-, interferon-, and NF-κB–associated transcriptional responses^100,101^, with pathway-level changes that opposed those induced by IL-1β stimulation in an independent dataset^67^. Moving to hematopoiesis, PerturbGen outperformed network-based inference and permutation baselines in a newly generated CD34⁺ HSPC atlas, identifying age- and lineage-specific PIPs in an *in silico* perturbation atlas featuring 3,108 gene perturbations, inspired by prior large-scale Perturb-seq screens^9,15,16^ to perform *in silico* reconstruction and reveal modular regulatory architecture governing dynamic, age-dependent hematopoiesis^3,4,70,71,74^. Leveraging naturally occurring human genetic variants to model perturbations of the same genes^102–104^, we were able to align hematological trait–associated genes with PerturbGen-defined PIPs and predict transcriptional alterations in a monogenic blood disorder. Finally, we went beyond an iterative trial-and-error approach, leveraging an *in silico* perturbation atlas of human iPSC-derived skin organoids to highlight a *GSK3B*/Wnt axis as a candidate for fibroblast maturation in skin organoids. This prediction aligns with multifaceted roles for Wnt signaling in murine dermal development^90,105–109^, and extends prior work showing that experimental activation of Wnt signaling via the GSK3β inhibitor, CHIR99021, increases skin organoid volume^98^, leveraging PerturbGen to demonstrate a program-level stromal shift towards *in vivo* cell states. Together, this demonstrates how trajectory-aware perturbation modeling can connect early signaling perturbations to later transcriptional outcomes, complementing within-state perturbation predictors that do not explicitly model propagation across state transitions^17,18,20,21^.

As with any trajectory-aware computational framework, PerturbGen’s has limitations. When intermediate time points or developmental stages are sparsely sampled, when transitions are poorly resolved, or when there is substantial heterogeneity within a state label, the model has less information to learn faithful state-to-state mappings and longer-range forecasting becomes increasingly challenging. Although PerturbGen currently models perturbations primarily at the transcriptional level using ranked tokenized gene expression, this approach does not explicitly represent post-transcriptional regulation, protein activity, chromatin state or cell-cell communication, which can be central to fate decisions and tissue organization. These limitations are particularly relevant for contexts where phenotypes are mediated by signaling dynamics, microenvironmental feedback or spatially structured interactions. More broadly, pre-training on a large and diverse corpus promotes generalization, but performance may be biased toward well-represented tissues, states and transitions. Incorporating richer structured metadata, such as tissue identity, spatial context and experimental covariates, could improve robustness and help disentangle biological variation from batch or sampling effects. Nevertheless, *in silico* gene screening allows comparison of predicted downstream effects across multiple unfolding cellular trajectories at a scale that would be technically infeasible using large-scale Perturb-seq screens^9,15,16^. Leveraging the predictive ability of PerturbGen, this approach has a myriad of potential applications. For example, PerturbGen can be applied to other complex and lineage-resolved cellular landscapes beyond hematopoiesis, spanning multiple experimental conditions and patient cohorts. In this context, *in silico* perturbations could provide a complementary means to model rare diseases that would otherwise remain challenging to study experimentally. Additionally, the opportunity to prioritize perturbations enables compensation for missing cues in current *in vitro* systems, such as organoids, identifying interventions that shift cellular compartments toward desired *in vivo* reference states and increasing the relevance of these systems for disease mechanism studies and therapeutic discovery.

In summary, PerturbGen extends modeling of gene perturbations from static to dynamic cellular systems, resolving lineage-specific programs, resolving rare human phenotypes and informing *in vitro* systems. As single-cell atlases expand across tissues, developmental windows and modalities^13,110^, PerturbGen provides a route to add perturbation-aware layers on top of these reference maps, complementing descriptive resources by enabling predictive and *in silico* hypotheses generation. We envision PerturbGen enabling the creation of *in silico*, trajectory-aware perturbation atlases and virtual cells across diverse biological scenarios, supporting optimization of human disease models and prioritization of candidate molecular interventions that redirect cells towards desired states and accelerate therapeutic discovery.

## Data Availability

All downstream differential expression predictions from *in silico* perturbations and perturbation atlases are available through the PerturbGen web portal at https://cellatlas.io/perturbgen.

## Code Availability

PerturbGen is available as a Python package, maintained at https://github.com/Lotfollahi-lab/Perturbgen/. All code to reproduce experiments, including benchmarking and analyses, is available at https://github.com/Lotfollahi-lab/Perturbgen-reproducibility.

## Acknowledgments

This project was conceived and funded by Open Targets, under OTAR 3090. We acknowledge core funding from Wellcome (WT220540/Z/20/A). The LPS work was funded by an MRC Large Collaboration Grant - Theme 2 AMR initiative: Optimising innate host defence to combatantimicrobial resistance (SHIELD consortium, MR/N02995X/1). M.H. is funded by Wellcome (WT107931/Z/15/Z) and the CIFAR MacMillan Multiscale Human program. T.I. was supported by the Funai Foundation for Information Technology and the Honjo International Scholarship Foundation. Work in the Gottgens laboratory was supported by Wellcome Trust [206328/Z/17/Z, 215116/Z/18/Z, 221052/C/20/Z], Medical Research Council [MR/W031663/1 and MR/V005502/1], Blood Cancer United [7035-24], Blood Cancer UK [18002] and Aging Biology Foundation. This research was funded in whole, or in part, by the Wellcome Trust [203151/Z/16/Z, 203151/A/16/Z, 226795/Z/22/Z] and the UKRI Medical Research Council [MC_PC_17230]. K.H.M.S was funded by Wellcome Trust [204017/Z/16/Z]. E.L. was funded by a Sir Henry Dale fellowship from Wellcome/Royal Society (107630/Z/15/Z). Research in EL’s laboratory was/is funded by Biotechnology and Biological Sciences Research Council (BB/P002293/1) and by Wellcome (215116/Z/18/Z and 309075/Z/24/Z). N.M. was supported by the Japan Society for the Promotion of Science Short-term Postdoctoral Fellowship and a Deutsche Forschungsgemeinschaft (DFG) Research Fellowship (ME 5209/1-1). D.J. is supported by a Wellcome Trust Accelerator Award (314710/Z/24/Z) and the NIHR Biomedical Research Centre at Great Ormond Street Hospital for Children NHS Foundation Trust and University College London. S.W. is funded by the Royal Society (CDF/R1/241008), and supported by the NIHR and Newcastle Biomedical Research Centre. J.S. is a National Institute for Health and Care Research (NIHR) Senior Investigator. The views expressed in this article are those of the author(s) and not necessarily those of the NIHR, or the Department of Health and Social Care. We thank Olivier Bakker for discussions on statistical evaluation and GWAS analyses, and Vicki Moignard for guidance in the interpretation of hematopoietic findings. We thank David Dixon and James Fletcher for processing the IV LPS samples for the Haniffa lab. We also thank members of the Lotfollahi and Trynka laboratories for discussions and feedback. This research was supported by the Cambridge BRC Cell Phenotyping Hub. We thank CRUK CI Genomics core for processing all Cambridge libraries/sequencing. For the purpose of open access, the author has applied a CC BY public copyright licence to any Author Accepted Manuscript version arising from this submission.

## Author Contributions

M.L.; B.G.; N.K.W.; and M.H.; conceptualized the study. K.CH.L.; led the project, with M.L.; leading manuscript preparation, A.V.; led overall model development, A.V.; and K.CH.L.; conceived and developed the model with feedback from M.L.; D.V. A.V.;K.CH.L;. D.V; M.P; A.H; and H.A; managed package and software development. M.M.; A.V.; K.CH.L.; A.M.F.; T.I.; and M.L.; designed the experiments. K.CH.L.; A.M.F.; and D.V.; performed the benchmarking, data simulation and ablation experiments with feedback from M.L.; and A.V.;. A.M.F.; curated pre-train data corpus with feedback from A.V;. A.V.; performed model pre-training, K.CH.L.; A.M.F.; and T.I. analyzed the data, with feedback from A.V.; G.T.; N.K.W; A.F.; E.G.; F.T.; M.S.V.; E.L.; and L.J.;. D.H.; and D.B.H.; designed and developed the web portal. Blood related CITE-seq datasets were generated by R.A.B.; N.M.; E.S.; D.I.; and N.K.W.; E.L. provided resources and supervised the generation of the blood CITE-Seq datasets; L.M. performed experiments. Computational analysis and QC of CITE-Seq datasets was performed by T.I.; M.Q.L.; M.S.V.; K.H.M.S.; and R.H.; Skin organoid datasets were generated by A.F.; F.T,; and C.A.;. K.CH.L.; A.M.F.; T.I.; D.J.; A.V.; M.L.; B.G.; N.K.W.; and M.H. wrote the paper. M.L.; B.G.; N.K.W.; G.T.; and M.H supervised the work. All authors reviewed the paper.

### Competing Interests

M.L. has equity interests in Relation Therapeutics, is a scientific co-founder and part-time employee of AI VIVO, and serves on the scientific advisory board of Novo Nordisk.

## Methods

### PerturbGen

PerturbGen is an encoder–decoder transformer^48,111^ that predicts downstream cellular gene expression states, enabling extrapolation along temporal or developmental trajectories, and inference of *in silico* perturbation effects on subsequent cell states. In this framework, the baseline condition is treated as the source sequence, intermediate conditions provide contextual information, and the downstream or perturbed state is treated as the target sequence. For time-resolved datasets, the source corresponds to a 0 h control state, the context consists of additional time points along the trajectory, and the target corresponds to a selected time point of interest, enabling either interpolation or extrapolation depending on target selection. For developmental datasets, the source represents a baseline or undifferentiated state, the context consists of additional stages along the differentiation trajectory, and the target corresponds to a selected stage of interest.

### Data pre-processing

#### Tokenizer

Let 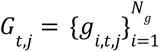 denote the gene expression profile of a cell 𝑗 ∈ {1, …, 𝐾} under condition 𝑡, where 𝑁 _𝑔_ is the total number of genes per cell, 𝐾 is the number of cells, and 𝑡 ∈ {0, …, 𝑇} represents discrete conditions (e.g., time points or stages). For simplicity, we omit the subscript 𝑗 in the following formulations, unless required for clarity.

A tokenizer 𝑟(·) converts continuous gene expression profiles into discrete gene tokens. For a cell in condition 𝑡, the resulting token sequence is

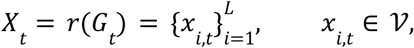

where 𝑥_𝑖,𝑡_ denotes the token corresponding to gene 𝑖 at condition 𝑡 and 𝐿 is the sequence length per cell (shorter sequences are padded to length 𝐿 using a token). The token vocabulary is defined as 𝒱 = 𝒱_𝑠𝑝𝑒𝑐𝑖𝑎𝑙_ ∪ 𝒱_𝑔𝑒𝑛𝑒_. We use |𝒱_𝑠𝑝𝑒𝑐𝑖𝑎𝑙_ | = 𝑁_e_ (e.g., [𝑀𝐴𝑆𝐾] and [𝑃𝐴𝐷]) and |𝒱 _𝑔𝑒𝑛𝑒_ | = 𝑁_g_ (gene tokens), resulting in |𝑉| = 𝑁_e_ + 𝑁_g_; thus 𝑋_t_∈ 𝒱 ^𝐿^ .

Gene tokens were derived using the Geneformer tokenizer^24^, with expression values normalized by gene-specific non-zero medians computed from our pre-training corpus. Expressed genes within each cell were ranked by normalized expression values, and the resulting rank order determined token positions. Tokenization and normalization were performed once across the full dataset prior to splitting into discrete conditions for downstream training.

### Cell pairing

ScRNA-seq does not allow individual cells to be profiled across multiple temporal or developmental stages. Because encoder-decoder transformers require explicit input-output pairs to learn conditional state transitions, we constructed pseudo-pairings between cells sampled at successive stages, forming ordered sequences spanning from an initial state to downstream cell states. When informative covariates are available (e.g., cell type or tissue), pairings are restricted within covariate strata to preserve biological consistency; otherwise, pairs are sampled at random within each successive stage. For developmental datasets with lineage annotations, pairings follow differentiation trees. To accommodate branching trajectories, a single source cell may be paired with multiple downstream target cells. Dataset-specific pairing strategies are summarized in Supplementary Table 15 and 18.

### Model Training

PerturbGen uses an encoder–decoder transformer architecture^48,111^ designed to predict downstream cellular gene expression states along temporal or developmental trajectories, as well as in response to perturbations. The model is trained in two phases: an encoder-only pre-training stage that learns general gene–gene relationships from large-scale single-cell transcriptomic data, followed by task-specific decoder training, in which downstream states are predicted using contextual information from intermediate conditions.

### Pre-training

The PerturbGen foundation model is pre-trained by training the transformer encoder on a large reference corpus of approximately 107 million single-cell transcriptomes (Fig. 1c,d). During this stage, only the encoder is trained using a masked language modeling objective. A subset of gene tokens is masked and predicted from the remaining context, i.e., unmasked tokens within each cell. Pre-training is conducted without cell state or temporal conditioning, enabling the encoder to learn general gene–gene relationships.

### Stage-specific decoder training

During task-specific training, the pre-trained encoder is kept fixed and its learned representations are reused. A randomly initialized decoder is optimized in two sequential stages. First, masked gene tokens are predicted for a randomly selected target condition 𝑡′, conditioned on unmasked tokens within the same cell and contextual information from intermediate time points or stages. We refer to this procedure as *decoder training*. Second, a cell embedding defined as the average of the gene token representations is used to reconstruct gene expression counts, providing additional training signal at the cell level.

### Model representations

Each gene token is mapped to a learnable 𝑑-dimensional embedding, randomly initialized and optimized during training. For a cell at condition 𝑡 with token sequence 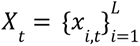, the corresponding embeddings are 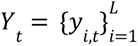, where 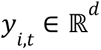. The cell embedding 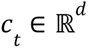 is computed as the mean of the token embeddings, excluding [𝑃𝐴𝐷] and [𝑀𝐴𝑆𝐾] tokens.

As self-attention mechanisms are permutation-invariant, we introduce positional encodings to incorporate sequence order into the model. Specifically, we use two positional encodings: a learnable embedding to represent the rank-defined ordering of gene tokens within each cell, and a sinusoidal encoding to represent temporal or developmental stage. This combination empirically yielded the best performance in our setting.

Formally, we add positional encodings

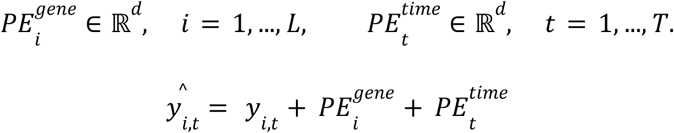

to the corresponding token embeddings before transformer processing. We denote the resulting positionally encoded sequence by 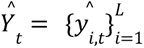, which is provided as input to the transformer.

### Masking strategy

During training, we sample a subset of token positions with probability β according to a masking scheduler γ. Using the same scheduler during inference aligns the masking ratio distribution across both phases, reducing distribution shift and leading to more stable and consistent generation. At each update, the masking ratio is determined by randomly sampling from the scheduler and applied in a single-step masking operation, following the MaskGIT^112^ training protocol. Padding tokens and other special tokens are excluded from masking. The mask index set is denoted by 𝑆 ⊂ {1, …, 𝐿}, |𝑆| = 𝑀, where 𝑀 represents the number of sampled tokens, and tokens in 𝑆 are replaced with [𝑀𝐴𝑆𝐾]. The resulting masked sequence is

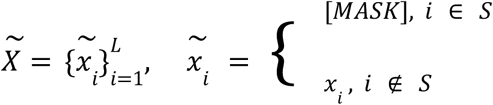

which serves as input to the model.

Ground-truth tokens at the masked positions are collected as

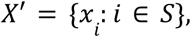

and are used as labels for the self-supervision task.

During pre-training, masking is applied to individual cell sequences without conditioning on temporal or cell state context. In contrast, during decoder training, if no target time step is specified, we randomly sample a target condition 𝑡′ from the set of available prediction cell states at each update step. Masking is then applied to the cells corresponding to 𝑡′. The model predicts masked tokens conditioned on the unmasked tokens of these cells via self-attention and on contextual embeddings from the remaining cell states via cross-attention.

### Context formation

During decoder training, representations from intermediate states are used to condition predictions for the target condition 𝑡′. The encoder 𝑓_δ_ processes the source condition 𝑡_0_ to produce a contextualized embedding 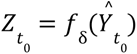. For each intermediate state 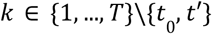, embeddings 𝑌_𝑘_ are processed by the decoder ℎ_θ_ in a forward-only manner to produce contextual embeddings 𝑍_𝑘_ . By projecting intermediate states through the decoder, we encourage it to learn temporally structured representations, enabling conditioning on cell state progression during target prediction. These embeddings are accumulated in a fixed, stage-ordered sequence to form a prefix of contextual information. The context used to predict the target condition 𝑡′ is then constructed as

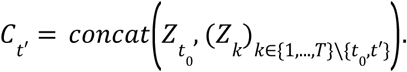

Context embeddings are computed without gradient flow, allowing information from intermediate states to condition prediction at 𝑡′ via cross-attention while preventing gradients from the target prediction from updating the context representations.

### Training objective

PerturbGen is trained using a masked token prediction objective that is shared between pre-training and decoder training but differs in how conditioning information is provided. During pre-training, masked gene tokens are predicted from individual cell sequences without contextual conditioning. During decoder training, the same objective is applied to a target condition 𝑡′, with prediction conditioned on contextual information from intermediate states.

Formally, for a target condition 𝑡′, the masked input sequence 𝑋_𝑡′_ is embedded as 𝑌_𝑡′_ and provided to the decoder as queries, while the context 𝐶_𝑡′_ is supplied as keys and values in cross-attention. The same decoder ℎ_θ_ is used to process both intermediate states and the masked target sequence, sharing parameters across all conditions. The model predicts the original tokens at masked positions 𝑆_𝑡′_ using a cross-entropy loss

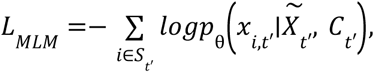

where

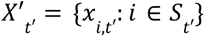 are the ground-truth tokens. In practice, one target condition 𝑡′ is sampled per update, and the rest serve as context, though in expectation, all conditions are covered during training.

### Gene expression decoder

Given a cell-level embedding 𝑐_𝑡_ ∈ ℝ*^d^* derived from mean-pooled token embeddings at condition 𝑡, gene expression counts 𝐺_𝑡_ are predicted using a lightweight count decoder adapted from scVI^113^. The decoder consists of a small multilayer perceptron that projects the cell embedding to a latent representation, followed by an L2 normalization layer to control embedding scale and stabilize count prediction across conditions. The normalized representation is then passed to output heads parameterizing a zero-inflated negative binomial (ZINB) distribution, which accounts for technical dropout and overdispersion in scRNA-seq data. The reconstruction loss is given by the negative log-likelihood of the observed counts under the ZINB model.

### Architectural and implementation

PerturbGen follows an encoder–decoder transformer architecture comprising a pre-trained twelve-layer transformer encoder and a condition-aware decoder. The decoder alternates self-attention and cross-attention layers to condition predictions on contextual information from intermediate states. Decoder depth is adjusted based on the task-specific dataset. Feedforward layers use standard transformer components, including layer normalization and GELU^114^ activations. Training and inference were optimized using PyTorch’s scaled dot-product attention (SDPA) implementation and mixed-precision computation to enable scalable training. We use Adam optimizer^115^. We use NVIDIA A100 80 GB and H100 80 GB for all the experiments. Full architectural specifications and hyperparameter settings are provided in the Supplementary Table 11.

### Model inference

#### Perturbation prediction

We modelled the effects of single-gene perturbations introduced at a source condition and predicted their transcriptional consequences in downstream target conditions. Perturbation simulation follows the strategy introduced in Geneformer, enabled by the shared ranked tokenization scheme. Gene knockdown or knockout perturbations are simulated by masking, padding, or removing the corresponding gene token, whereas gene overexpression is simulated by repositioning the token to the highest rank within the sequence. PerturbGen differs from Geneformer in that perturbations are applied at the source condition, and their effects are propagated to one or more downstream target conditions.

Predicted perturbation effects can be evaluated at the level of latent representations or reconstructed gene expression using the count decoder. For each perturbation, two forward passes were performed (control and perturbed). Perturbation effects were quantified using (i) changes in cell embeddings measured by cosine similarity and (ii) predicted transcript-level responses derived from the reconstructed gene expression profiles. Differential expression analysis was performed using sc.tl.rank_genes() from Scanpy^116^ to estimate log_2_ fold changes and perturbation directionality.

### Cell sequence generation

Gene expression is not inherently sequential; instead, transcriptional states arise from coordinated and nonlinear gene regulatory interactions. We therefore employ a non-autoregressive iterative decoding strategy inspired by MaskGIT^112^, which predicts masked tokens in parallel while leveraging bidirectional context, analogous to BERT^117^. Generation is initialized from a fully masked sequence, in which all non-padded tokens are replaced with a [MASK] token, denoted as 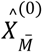. At iteration 𝑟 ∈ {1, …, 𝑅}, the masking scheduler γ determines the number of tokens that remain masked:

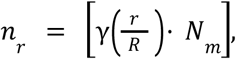

where 𝑁_𝑚_ is the total number of gene tokens to be generated and γ(.) is a masking scheduling function (e.g., cosine or power decay). As the iteration index increases, the number of mask tokens 𝑛 _𝑟_ decreases. At iteration 𝑟, the model predicts categorical probability distributions over the vocabulary for all masked positions in parallel, denoted 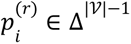 conditioned on the bidirectional context provided by the unmasked tokens. Tokens are sampled from 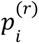 with temperature annealing to control generation diversity.

Because each gene token may occur at most once per sequence, tokens selected in previous iterations are excluded from subsequent sampling. The iterative prediction procedure is repeated until iteration 𝑅, at which point all gene tokens have been generated.

### Pre-training data

#### Data curation

We assembled a large-scale pre-training corpus comprising 107 million (106,967,684) human single-cell transcriptomes spanning a broad range of tissues and disease states, derived from a combination of publicly available and private datasets (Fig. 1c,d; Supplementary Table 1).

Publicly available datasets containing raw count matrices were obtained primarily from the National Center for Biotechnology Information (NCBI) Gene Expression Omnibus (GEO), the CellxGene^118^ data release dated 2025.06.01, and the scBaseCount^119^ pre-training corpus (release 2025.06.01). For datasets obtained from scBaseCount, studies were restricted to droplet-based 10x Genomics 3′ sequencing data to ensure consistency across samples. Additional datasets sourced from repositories other than CellxGene are detailed in Supplementary Table 1.

To prevent the inclusion of duplicated cells, we preferentially used primary data from CellxGene when available. For datasets obtained from other resources, metadata and study-level information were manually reviewed to identify and exclude potential overlaps. In total, 9,912 unique h5ad files were included for model pre-training.

Each dataset was processed independently using a uniform preprocessing and quality-control pipeline. Only raw count expression matrices were used, and datasets lacking raw count data were excluded. Gene annotations were harmonized using Ensembl identifiers, and gene filtering was applied to retain high-confidence protein-coding genes while excluding non-coding, LINC and MIR genes.

Per-cell quality-control metrics were computed for each dataset using Scanpy, including total UMI counts, number of detected genes per cell (gene complexity), fraction of counts attributed to the top 20 expressed genes, and percentage of mitochondrial reads. Extreme values were identified using a median absolute deviation (MAD)-based approach. Cells were excluded if they deviated by more than five MADs from the median for log-transformed total counts, log-transformed number of detected genes, or the percentage of counts in the top 20 genes. Mitochondrial outliers were defined as cells deviating by more than four MADs from the median mitochondrial fraction or exhibiting >20% mitochondrial reads. Genes detected in fewer than five cells were removed. All quality-control procedures were performed independently for each dataset prior to model pre-training.

### Datasets

As part of this study, we generated and made available three previously unpublished datasets: in vivo lipopolysaccharide (LPS) challenge, human haematopoiesis, and skin organoid development.

### *In vivo* LPS challenge

#### Sample acquisition

For the intravenous (IV)-LPS samples, ethical approval was granted by a Research Ethics Committee (17/YH/0021). Healthy volunteers were screened against the inclusion criteria in the study protocol (Supplementary File 1) and those that passed were recruited and provided informed, written consent. The LPS (Clinical Center Reference Endotoxin, 94332B1) was obtained from the National Institute of Health and was injected intravenously as a bolus dose of 2 ng kg^−1^. A baseline blood sample in ethylenediaminetetraacetic acid (EDTA) tubes was taken before the LPS administration and then again at the following time points: 90 minutes, 6 hours and 10 hours after administration.

### Sample processing

PBMCs were then harvested from the blood samples using Lymphoprep (StemCell Technologies) according to the manufacturer’s protocol, washed with Dulbecco’s PBS (Sigma) and frozen in 90% (vol/vol) heat-inactivated FCS (Gibco) and 10% (vol/vol) DMSO (Sigma-Aldrich).

### CITE-seq experiments

On the day of the experiment, cells were thawed for 1 minute and washed with PBS supplemented with 2% (vol/vol) FCS and 2 mM EDTA (Sigma), before centrifuging for 5 minutes at 500g. Cells were then resuspended and filtered using a 30 μm filter before counting. Dead cells were removed using the MACSdead cell removal kit (Miltenyi Biotech) according to the manufacturer’s protocol. 200,000 live cells from each donor were stained with Human TruStain FcX Fc Blocking Reagent (BioLegend, 422302) for 10 min at room temperature. A custom Total-Seq-C panel (BioLegend, 99813; Supplementary Table 2) was then used to stain the cells for 30 min at 4 °C. Cells were washed twice with PBS supplemented with 2% (vol/vol) FCS and 2 mM EDTA (Sigma) before resuspending in PBS. Approximately 20,000–30,000 cells per sample were processed using Chromium NextGEM Single Cell V(D)J Reagent kits v1.1 with Feature Barcoding technology for Cell-Surface Protein (10x Genomics) according to the manufacturer’s protocol. Libraries were then sequenced using Illumina sequencing technology (NovaSeq 6000) to achieve a minimum of 50,000 raw reads per cell.

Data from time points 90 minutes and 10 hours have been published as part of our previous study^54^.

### Preprocessing and annotation

LPS scRNA-seq data were processed using Scanpy. Low-quality cells with mitochondrial transcript content > 8% were excluded, and genes expressed in fewer than 20 cells were removed. Data were normalized to a constant total count per cell (10,000 reads) using the `normalize_total` function and log-transformed. Cell type annotation was performed using the automated CellTypist pipeline^120^ with the human PBMC reference model. Dimensionality reduction was performed by PCA using 50 principal components. A *k*-nearest neighbor graph was constructed using the `neighbors` function, followed by UMAP for visualization.

### Human hematopoiesis

#### Tissue acquisition

Human developmental tissues (yolk sac (YS), fetal liver (FL) and fetal bone marrow (BM)) were obtained from the Human Developmental Biology Resource (HDBR), following elective termination of pregnancy, with written informed consent and approval from the Newcastle and North Tyneside NHS Health Authority Joint Ethics Committee (18/NE/0290 and 08/H0906/21+5). The HDBR is regulated by the UK Human Tissue Authority (HTA; https://www.hta.gov.uk/) and operates in accordance with the relevant HTA Codes of Practice. Cord blood (CB) samples were obtained from Cambridge Blood Stem Cell Biobank, in accordance with regulated procedures approved by the relevant Research and Ethics Committees (18/EE/0199). Pediatric BM samples were collected after written informed consent, in accordance with the Declaration of Helsinki under a study approved by the National Research Ethics Committee (12/LO/0426). Healthy donor pediatric BM was collected from sibling donors. Young adult tissues were purchased from Stem Cell Technologies. Aged adult tissues were obtained from Newcastle Hospitals NHS Foundation Trust, following hip replacement surgery, with written and informed consent and approval from the Newcastle and North Tyneside NHS Health Authority Joint Ethics Committee (19/LO/0389).

### Tissue processing

Adherent material was removed, and the fetal femur was cut into small pieces before grinding with a pestle and mortar. A flow buffer (PBS containing 5% (v/v) FBS (Gibco) and 2 mM EDTA (Sigma)) was added to reduce clumping. The suspension was filtered with a 70-μm filter and red cells were lysed with 1× RBC lysis buffer (eBioscience) according to the manufacturer’s instructions.

### CITE-seq experiments

Cryopreserved YS (n = 2; 6–7 PCW), FL (n = 9; 6–17 PCW), fetal BM (n = 3; 14–17 PCW), CB(n = 4), pediatric BM (n = 4; 1–16 years old) and adult BM cells (n = 9; 29–91 years old) (Supplementary Table 3) were thawed on the day of experiment and added to pre-warmed RF-10 (RPMI (Sigma-Aldrich) supplemented with 10% (v/v) heat-inactivated FBS (Gibco), 100 U ml^−1^ penicillin (Sigma-Aldrich), 0.1 mg ml^−1^ streptomycin (Sigma-Aldrich), and 2 mM L-glutamine (Sigma-Aldrich). CD34^+^ selection was performed on adult BMcells using CD34 MicroBead Kit (Miltenyi Biotec) and manual separation on LS columns. Cells were manually counted and pooled if cell numbers were low (pools noted in the sequencing index columns; Supplementary Table 3). After incubation with Fc receptor blocking reagent (BioLegend), cells were labelled with CD34 APC/Cy-7 (Supplementary Table 4) for 10 min in the dark and on ice. During the incubation, the CITE-seq^69^ antibody cocktail vial was centrifuged at 14,000g for 1 min then reconstituted with flow buffer. The vial was incubated for 5 min at room temperature then centrifuged at 14,000g for 10 min at 4 °C. The CITE-seq antibody cocktail (Supplementary Table 4) was then added to the cells along with a competition antibody mix (Supplementary Table 5) and incubated for 30 min in the dark and on ice. The stained cells were then washed with flow buffer before resuspension in flow buffer supplemented with 50 μg ml^−1^ 7-AAD (Thermo Fisher Scientific).

Live, single CD34^+^ cells were sorted by FACS into 500 μl PBS in pre-chilled FACS tubes coated with FBS until the sample was exhausted. Samples were sorted at the NIHR Cambridge BRC Cell Phenotyping Hub using the BD FACSAria Fusion and FACSAria III sorters. Sorted cells were then centrifuged at 500g for 5 min before manual counting. Cells were then submitted to the CRUK CI Genomics Core Facility for 10x Chromium loading, library preparation and sequencing. Single cell 3′ v3 (10x Genomics) kits were used and gene expression and cell-surface protein libraries were generated as per the manufacturer’s protocols. Libraries were sequenced using an Illumina Novaseq platform. The YS, FL, fetal BM, CB and PBM samples have been published as part of our previous studies^70,74,121^.

### Alignment and preprocessing of CITE-seq data

CITE-seq transcriptomic data were aligned to the GRCh38 genome and quantified using the Cell Ranger pipeline (v4.0.0). Cell-associated barcodes and background-associated barcodes were determined using the EmptyDrops method^122^ implemented in the Cell Ranger pipeline, and the background-associated barcodes were excluded. Pooled samples were demultiplexed with Souporcell^123^ and Vireo^124^, where individual donor-derived cells were first clustered using Souporcell and donor information was matched using Vireo based on paired SNP array data (UK Biobank Axiom Array, Applied Biosystems, performed by Cambridge Genomic Services, University of Cambridge) obtained for each pooled donor. Subsequent data analysis was performed using Scanpy^116^. Multiplets were removed using Scrublet^125^ based on the threshold of the median plus three times the median absolute deviation scrublet score, as previously described^71^. Cell libraries with less than 1,000 detected genes or with mitochondrial gene expression exceeding 10% of unique molecular identifier (UMI) counts were removed.

Cell surface protein data were quantified using CITE-seq-Count^126^ (v1.4.3) with the option ‘-cells 200000’ to return sufficient background droplets for the subsequent denoising analysis. Cells with more than 140 detected proteins or with total protein counts less than the median minus 0.8 times the median absolute deviation were further removed. Data for the remaining 102,744 cells were denoised and normalized using DSB^127^. The DSB-normalized protein expression matrix was used to compute 50 principal components (PCs), and the PCs were adjusted for batch effects between the individual samples using Harmony^128^. The 50 batch-corrected PCs were used to identify 12 nearest neighbors and to compute clusters using the Scanpy functions `neighbors` and `leiden`, and a B/myeloid doublet cluster expressing both CD19 and CD33 was removed.

After these transcriptome- and protein-based quality control steps, the gene expression count matrix for the remaining 101,477 cells was log-normalized, and 2,000 highly variable genes were identified. Cell cycle scores were computed and subsequently regressed out using a previously published list of cell cycle-associated genes^129^ and the Scanpy functions `score_genes_cell_cycle` and `regress_out`.

### Dimensional reduction and annotation

The pre-processed gene expression matrix was scaled and used to compute 50 PCs. Protein expression-based PCs were recomputed for the remaining 101,477 cells. The transcriptome-and protein-based PCs were separately adjusted for batch effects between the individual samples using Harmony^128^. A multimodal neighbor graph was constructed based on the 50 transcriptome-based and 50 protein-based batch-corrected PCs using the weighted nearest neighbor (WNN) algorithm^130^. The WNN graph was then used to compute clusters and the UMAP embedding using the Scanpy functions `leiden` and `umap`, respectively. Data annotation was performed manually using marker genes and proteins identified through literature search (Supplementary Table 6). Particularly, the following populations were annotated based on their surface protein phenotypes: LT-HSCs (CD34^hi^ CD38^neg–lo^ CD90^hi^ CD45RA^neg–lo^ CD49f^hi^)^131^, ST-HSCs (CD34^hi^ CD38^neg–lo^ CD90^hi^ CD45RA^neg–lo^ CD49f^neg–lo^)^131^, MPPs (CD34^hi^ CD38^neg–lo^ CD90^neg–lo^ CD45RA^neg–lo^)^132^, LMPPs (CD34^hi^ CD38^neg–lo^ CD90^neg–lo^ CD45RA^hi^)^133^, CLPs (CD34^hi^ CD38^hi^ CD10^hi^ Lineage^neg–lo^)^134^ and GMPs (CD34^hi^ CD38^hi^ CD45RA^hi^ CD10^neg–lo^)^135^ (Supplementary Fig. 10b and 11). CD34^+^ stromal cell populations, including *CDH5*^+^ endothelial cells^136^, *COL1A1*^+^ *DCN*^+^ fibroblasts^137^ and *ALB*^+^ *AFP*^+^ hepatocytes^138^, were also identified and excluded from our hematopoietic landscape. For the remaining 98,266 cells, the final UMAP embedding was recomputed using the `umap` function.

### Published skin organoid

We used preprocessed scRNA data from Gopee *et al*^6^, comprising samples collected at days 29, 48, 85, and 133 of skin organoid differentiation, which were merged with day 6 samples obtained from Lee *et al*^90^. The merged dataset was inspected to ensure consistency in gene annotation and metadata structure across studies prior to subsequent computational analyses.

### GSK3β-inhibited skin organoid

#### Skin organoid differentiation and perturbation experiments

The hair-bearing skin organoids were generated from Kolf2.1S human induced pluripotent stem cells (iPSCs) as previously described, with minor modifications to the protocol for WNT signaling activation^88^. For activation of the canonical WNT signaling pathway, on day 6 of the culture, the organoids were treated with 3 μM Wnt agonist (CHIR99021; Tocris). At the target time points, at least 3 organoids were selected, pooled and dissociated as follows. The organoids were incubated on a rotator (65 rpm) at 37°C for 45 min in pre-warmed RPMI supplemented with 10% FBS and collagenase IV, with intermittent pipetting every 10 min. Following this step, the tissue was filtered through a 70 μm filter and the cells were collected. Any undissociated tissue left over on the filter, was then incubated with TrypLE™ Express Enzyme (1X), phenol red (Gibco) for 10-20 min on the rotator (65 rpm) at 37°C. Following the two-step digestion, the cells were washed in PBS supplemented with 3% BSA and then resuspended in PBS supplemented with 0.4% BSA. ScRNA-seq was performed as per manufacturer’s instruction using v3.1 chemistry (10x Genomics user guide CG000204 Rev D). The libraries were sequenced on Illumina Novaseq platform.

### Data preprocessing and QC

Per-cell QC metrics were computed, and cells with total UMI counts between 7,000 and 50,000 and mitochondrial transcript content <20% were retained, excluding low-quality or stressed cells. Putative doublets were identified using Scrublet^125^ and excluded from downstream analyses. Normalization was performed using deconvolution-based size factor estimation implemented in Scanpy, followed by log-transformation of normalized counts. Library size distributions were inspected across time points and conditions. To correct the sequencing-depth imbalance observed at day 70, unperturbed cells were subsampled using bin-wise matching to align their total UMI count distribution with that of perturbed cells, minimizing potential confounding effects of library size on downstream comparisons.

The rebalanced dataset was modeled using scVI with default parameters on raw counts, conditioning on sample observation as categorical covariate and mitochondrial transcript fraction as continuous covariate. Top 1,200 highly variable genes were selected using the Seurat v3 method, specifying time point as a batch variable during gene selection. The resulting latent space was used to construct a neighborhood graph, from which Leiden clustering was performed and UMAP embeddings were computed. Cell type annotation was performed manually based on the expression of established canonical marker genes (Supplementary Table 9; Supplementary Fig. S20b).

### Benchmarking

#### Time point interpolation and extrapolation

We evaluated the generative performance of PerturbGen against established trajectory modeling approaches, including MIOFlow^41^, OT-CFM^49^ and Prescient^37^. Benchmarking followed the experimental design as previously described^36^, and OT-CFM was implemented using the authors’ publicly available repository. Models were assessed across three independent random initializations of model parameters, with dataset shuffling held constant across runs. Evaluation was performed on three time-resolved scRNA-seq datasets: *in vivo* LPS stimulation, T cell activation^139^ and embryoid body differentiation^140^.

For all datasets, 2,000 highly variable genes were selected using sc.pp.highly_variable_genes() from Scanpy. During training, all observed time points except the designated held-out target time point were provided to the model; the held-out time point was used exclusively for inference. PerturbGen was trained on tokenized gene expression representations, with upsampling applied as needed to balance cell numbers. A count decoder produced expression estimates for the complete set of 2,000 highly variable genes, ensuring comparability across methods.

Baseline methods were trained in a PC space derived from log-normalized training data retaining 50 PCs for MIOFlow and Prescient and 100 PCs for OT-CFM, following prior work. PCA was fit using training time points only to prevent data leakage. Predicted PC embeddings were inverse-transformed to gene expression space for evaluation.

Generative performance was quantified by comparing predicted and observed gene expression distributions at the held-out time point. Maximum Mean Discrepancy^141^ (MMD) was computed using a radial basis function (RBF) kernel with bandwidth parameters γ ∈ {2, 1, 0. 5, 0. 1, 0. 01, 0. 005} and 10,000 cells randomly sampled from predicted and observed distributions; the final value was averaged across kernel bandwidths. Earth Mover’s Distance^142^ (EMD; 1-Wasserstein distance) was computed using all available cells; when duplicate cells were present, mean expression values were used for evaluation. All metrics were calculated on log-normalized expression values. Hyperparameters for PerturbGen are provided in Supplementary Table 11. Baseline hyperparameters followed those reported in Zhang *et al.*^36^ and the original OT-CFM publication.

### Ablation analyses for time point prediction

We performed systematic ablation studies to evaluate the contribution of architectural components and MaskGIT-style iterative demasking strategies^112^ to time point prediction performance (Supplementary Fig. 2). Model performance was assessed using EMD and MMD.

### Sequence length ablation

We varied the number of masked tokens predicted per sequence in 25-token increments. Remaining tokens were padded to the maximum sequence length.

### Count prediction ablation

To assess the impact of probabilistic count modeling, we evaluated zero-inflated negative binomial (ZINB) sampling with varying numbers of samples during inference.

### Positional encoding ablation

Three encoding schemes were evaluated: (i) *sin_learnt*, combining sinusoidal time point encoding with learnable gene-rank embeddings; (ii) *pos_sin*, applying independent sinusoidal encodings to time points and gene ranks; and (iii) *comb_sin*, jointly encoding time points and gene ranks within a unified sinusoidal representation.

### Iterative demasking strategy

A standard top-*k* strategy was compared with a top-*k* probability-margin strategy^143^ that ranks positions based on the difference between the two most probable tokens, prioritizing positions with greater predictive certainty. Given its superior performance, the top-*k* probability-margin strategy was used by default.

### *In silico* perturbation prediction

Predicted perturbation-induced transcriptional changes were benchmarked against a previously published multi-omic pooled transcription factor (TF) CRISPR knockout dataset^75^. The dataset was used as provided in the original publication, and downstream analyses were conducted using log-normalized count matrix. The analysis was restricted to 9 of 19 TFs exhibiting >3 up-and down-regulated pathways (Supplementary Fig. 12b). Models were trained on an unperturbed human HSPC dataset, and TF loss was simulated *in silico* at inference. For the benchmark, masking of gene in the source sequence was used to induce *in silico* perturbation. For TFs profiled in both datasets, predicted gene-level expression changes were compared with experimentally observed effects to assess directional concordance.

PerturbGen was trained on the HSPC landscape. *In silico* perturbations were introduced in megakaryocyte–erythroid–mast cell progenitors (MEMPs), and downstream effects were predicted across differentiated lineages, including megakaryocytes, erythrocytes and basophil–eosinophil–mast cell populations.

Cluster-specific GRN were inferred from the HSPC scRNA-seq dataset (GSE96769) using CellOracle^28^, integrating orthogonal ATAC-seq data from mouse and human HSPCs to constrain TF–target interactions by chromatin accessibility and motif enrichment. The mouse GRN was obtained from the CellOracle tutorial resource. For the human GRN, single-cell ATAC-seq data (109,418 peaks × 2,210 cells) were binarized and processed with Monocle3 (LSI and UMAP). Cis-regulatory co-accessibility networks were inferred using Cicero (50 cells per bin), and distal peaks were linked to target genes by integrating TSS annotation (hg19) with co-accessible peaks, retaining connections with co-accessibility ≥0.8. Motif scanning was performed using CellOracle (hg19, FPR 0.02), and motif-supported TF–target pairs were used to construct the regulatory network. Default CellOracle parameters were applied unless otherwise specified. The resulting TF-peak associations were mapped to genes via peak-to-gene links derived from TSS annotation and co-accessibility, yielding the final human TF–target network used for GRN inference in CellOracle.

Perturbation simulations were performed on the full dataset and subsequently restricted to the same cell population (excluding MEP, early GMP and EoBasoMast precursor cells) used in PerturbGen to ensure comparability. To perturb TFs, expression of the target TF was set to zero in each cell, followed by iterative signal propagation through the inferred GRN to estimate downstream transcriptional responses. The number of propagation steps (n_propagation) was set to 3, as this value maximized agreement between predicted and observed perturbation-induced gene expression changes in held-out TF perturbations (Supplementary Fig. 13g, h).

Predicted perturbation effects were quantified as changes in gene expression relative to the unperturbed state. To quantify directional concordance, DEG analysis was performed in the experimental dataset using a Wilcoxon rank-sum test with Benjamini–Hochberg correction^144^ to define ground-truth differentially expressed genes (DEGs) (adjusted *P* value < 0.05) based on log_2_ fold-changes. Subsequent evaluation was restricted to these ground-truth DEGs. Balanced accuracy was computed as the proportion of genes for which the predicted and observed perturbation directions were concordant. Balanced accuracy was defined as the mean of sensitivity and specificity for predicting DEG directionality (up- versus down-regulated pathways). Performance was further evaluated using the macro F1 score, equally weighing up-and downregulated genes, as implemented in scikit-learn^145^. Predictions with zero change were treated as negative (no-effect) predictions in the classification analysis. Metrics were averaged across TF perturbations.

### Data Analyses

#### Gene embedding clustering and scoring

Gene embeddings were averaged across all cells within each time point, considering only cells in which the corresponding genes exhibited non-zero expression. For each gene, this resulted in three timepoint-specific embeddings, each represented as a 768-dimensional vector. Dimensionality reduction was performed using PCA with default Scanpy parameters. A *k*-nearest neighbor graph was subsequently constructed, and UMAP was applied for visualization. Leiden clustering was applied in order to identify gene programs.

The cytokine signaling gene program was scored using the score_genes function in Scanpy, which computes the average expression of the specified gene set for each cell individually.

### Pathway-level comparison of IL-1β stimulation and in silico *IL1B* knockout

Preprocessed cytokine screen data were obtained from the Parse Biosciences database^67^, and no additional preprocessing was applied. The LPS dataset was restricted to myeloid lineage cell types (CD14+ monocytes, CD16+ monocytes and DCs) exhibiting the strongest perturbation effects as identified by PerturbGen. Both the predicted *in silico IL1B* KO data and the cytokine stimulation dataset were restricted to the myeloid lineage and subset to an overlapping 1,990-gene panel to ensure comparability.

DEGs were identified using the Wilcoxon rank-sum test in Scanpy. In the experimental ground-truth dataset, IL-1β-stimulated cells were compared with PBS-treated controls. In the *in silico* perturbation, perturbed and unperturbed cells were compared at 6 h and 10 h. Genes with a Benjamini–Hochberg adjusted *P* value < 0.05 and absolute log_2_ fold-change > 0.2 were considered significant.

Pathway over-representation analysis (ORA) was performed using EnrichR via GSEApy^146^, querying the Reactome_Pathways_2024 gene set collection (Homo sapiens) with the shared 1,990 gene panel as background. Upregulated and downregulated gene sets were analyzed separately. Pathways with an adjusted *P* value < 0.05 and containing at least five genes were retained for downstream analysis.

For each pathway, constituent genes were extracted from the enriched IL-1β pathway sets and their log_2_ fold-changes in the *in silico IL1B* KO dataset at 6 hour and 10 hour were retrieved from differential expression analysis. Gene-level direction was assigned by the sign of the log_2_ fold-change. Pathway-level directionality was summarized by summing gene-level directional signs across all genes within each pathway, such that pathways with predominantly upregulated genes received positive scores and those with predominantly downregulated genes received negative scores.

To facilitate comparison across conditions, pathway scores were normalized separately within each condition by dividing by the maximum absolute pathway combined score observed in that condition, resulting in scaled values bounded between −1 and 1. This normalization was applied solely for visualization and does not alter the relative directionality within each condition.

Reverse concordance was defined as pathways exhibiting opposite overall directional trends between IL-1β stimulation and *in silico IL1B* KO.

### Quantitative weighted pathway-level reversal accuracy

To quantify reversal of IL-1β induced transcriptional programs by PerturbGen, we implemented a pathway-level weighted reversal framework. Differential expression analysis between IL-1β-treated and PBS control cells was computed in the Parse Biosciences dataset using a Wilcoxon rank-sum test. Genes with adjusted P < 0.05 and absolute log fold change > 0.5 were defined as the ground-truth (GT) IL-1β response and partitioned into upregulated and downregulated subsets according to direction of change.

ORA was performed using the Reactome Pathways (2024 release) gene sets. Enrichment was assessed on GT genes, restricting pathway sizes to 5–500 genes and applying multiple testing corrections (adjusted P < 0.05). Significant enrichment was observed only for the upregulated GT gene set, which defined biologically coherent pathways representing the IL-1β transcriptional program.

For *in silico IL1B* KO (6 h and 10 h), differential expression was computed between predicted perturbed and unperturbed conditions. Gene-level direction was defined by the sign of the log fold change. A gene was considered reversed if its predicted direction was opposite to the GT direction. Each gene’s contribution was weighted by the absolute magnitude of its GT log fold change, thereby assigning weights proportional to IL-1β response magnitude.

For each enriched pathway, we calculated the fraction of the total GT-weighted signal reversed by the model. Pathway-level reversal scores were then averaged across all significant pathways to obtain a global weighted reversal statistic for each condition.

Statistical significance was assessed using an empirical null distribution generated by randomly permuting predicted gene directions while preserving the set of evaluable genes. For each permutation, the pathway-level weighted reversal statistic was recomputed. Empirical *P* values were calculated as the proportion of permutations yielding reversal scores greater than or equal to the observed value.

### Perturbation atlas

During inference, *in silico* perturbations were exclusively introduced at the source state by modifying the token corresponding to the target gene. Specifically, the gene token was either replaced with a [𝑀𝐴𝑆𝐾] token, replaced with a [𝑃𝐴𝐷] token or removed entirely from the input sequence. Model performance was evaluated across these strategies, and the [𝑀𝐴𝑆𝐾] condition was selected for subsequent perturbation analyses based on predictive performance (Supplementary Fig. 13e, f).

For the HSPC dataset, model training was run on 5k highly variable genes. We further ensured that relevant transcription factors (TFs) are included during model training by incorporating TF expressed in >1000 cells (Supplementary Table 14). Perturbations were introduced in all intermediate cells (GMP, MEMP and LMPP) in which the gene exhibited non-zero expression. Together, they form the entire vocabulary size, a total of 5848 genes, to train PertubGen. For the skin dataset, model training was performed using 5,000 highly variable genes. To ensure adequate representation of developmental regulators, TFs implicated in development were additionally included in the training set (Supplementary Table 17). Perturbations were simulated in all day 6 cells in which the target gene exhibited non-zero expression. In total, these genes constituted the full model vocabulary, comprising 5,050 genes, which was used to train PertubGen.

To construct a perturbation atlas, cell embeddings were mean-aggregated across all cells in which the perturbed gene showed non-zero expression, either globally or within selected conditions. For the HSPC dataset, only perturbations achieving a predicted gene expression Pearson correlation coefficient greater than 0.90 were retained. For the skin dataset, a threshold of 0.60 was applied to account for temporally resolved perturbation responses.

Perturbation-induced programs (PIPs) were identified using Scanpy. A *k*-nearest neighbour graph was constructed from the aggregated embeddings, followed by Leiden clustering `leiden`. The number of neighbours and clustering resolution were selected by optimizing clustering quality, assessed using silhouette score and cluster granularity.

The final gene clusters were assessed for gene ontology enrichment using Metascape^147^, and gene program activity was computed using the Scanpy function `score_genes`. Gene programs were then annotated based on enriched gene ontology terms and age- and lineage-specific expression patterns (Supplementary Fig. 14a-f).

### HSPC differential abundance analysis

Differential abundance analysis was performed using the Python package MELD^148^, as implemented in our previously published PMCA pipeline^149^. Briefly, a cell similarity graph was constructed based on Euclidean distances in the 100-dimensional PC space (i.e., 50 RNA and 50 protein PCs), and a kernel density estimate (KDE) was computed for each sample. Differential abundance was then quantified by performing cell-wise L1 normalization of KDEs between individual samples. Young adult BM samples (29–50 years old) were used as the reference group and compared against each age group. The sample-associated relative likelihoods from all pairwise comparisons were averaged to obtain the age group-specific differential abundance landscapes.

### Permutation test for pathway analysis

Perturbation pathway-level effects were quantified in both experimental and *in silico* datasets using gene set enrichment analysis. For cross-dataset comparison, the union of expressed genes between the two datasets was used as the fixed ranked gene universe. Gene-level ranks were obtained using the Wilcoxon rank-sum test implemented in Scanpy. To mitigate stochasticity arising from rank ties, small random values (10^−12^) were added prior to ranking. Ranked gene lists were provided to GSEApy^146^, restricting analysis to gene sets containing 10-500 genes from Reactome (2024 release). Internal GSEA permutations were used solely to estimate enrichment statistics (normalized enrichment score (NES) and FDR) and were not used for cross-dataset significance testing.

Experimental GSEA results were treated as ground truth, with significant pathways defined by false discovery rate (FDR) q-value < 0.05 and an absolute NES > 1.0. The null hypothesis of the permutation test was that predicted gene rankings are independent of experimentally observed pathway directionality. Pathway direction was defined by the sign of the NES. For each permutation (B = 5,000), predicted gene ranks were randomly permuted across genes within the fixed gene universe, GSEA was recomputed, and pathway directionality and balanced accuracy were recalculated. The empirical one-sided *P* value was calculated as the proportion of permutations yielding a balanced accuracy greater than or equal to that observed for the PerturbGen predictions.

### Disease simulation in ETV6 thrombocytopenia patients

Transcriptomic responses to *in silico ETV6* KO in the HSPC perturbation atlas were compared with an independent scRNA-seq dataset of *ex vivo*-derived megakaryocytes from ETV6 thrombocytopenia patients^87^ (two controls and two patients, one harboring a missense and one a nonsense mutation). Analyses were restricted to megakaryocytes to assess disease-associated transcriptional effects within this lineage.

Data processing, genotype assignment and hashtag oligonucleotide (HTO) demultiplexing were performed as described in the original study, except that raw counts were log-normalized prior to analysis. Directional concordance of pathway effects between the *in silico* perturbation and patient data was assessed using the permutation framework described above, applied prior to all post-processing and cell-level filtering steps described below.

Souporcell genotype assignments were integrated with HTO demultiplexing to define donor identities and case–control labels. HTO singlets were assigned by donor identity. HTO-negative cells were retained only if they belonged to genotype clusters with minimal cross-donor mixing and showed strong support for singlet classification (ΔlogP = log_prob_singleton − log_prob_doublet > 2). Doublets and low-confidence cells were excluded.

Cells expressing fewer than 100 genes and genes detected in fewer than three cells were removed. To minimize contamination from myeloid populations, analyses were restricted to cells expressing *GP9* and *ITGA2B* (log-normalized count > 0) and lacking detectable expression of *LST1* and *TYROBP* (log-normalized count = 0).

Gene program activity was quantified using `score_genes`. Two predefined gene sets were evaluated: an MHC class II program (*CD74*, *HLA-DRA*, *HLA-DPA1*, *HLA-DMA*, *HLA-DRB1* and *CTSS*) and a megakaryocyte program (*PF4*, *PPBP*, *NFE2*, *VWF*, *TUBB1*, *ITGA2B*, *GP9* and *GP1BA*). Given the limited number of biological replicates (n = 2 per group), group differences are presented descriptively without formal statistical testing.

### GWAS trait analysis using OpenTargets

For the perturbed genes in the perturbation atlas, corresponding GWAS trait-gene associations were retrieved from the Open Targets Genetics^79^ integrated GraphQL endpoint (Version 25.09; accessed on 14 November 2025). GWAS-based associations were filtered to exclude low-confidence Locus-to-Gene (L2G) links (score < 0.25). Curated genetic disorder annotations were filtered independently to exclude entries with limited supporting evidence (ClinVar < 0.5 or PanelApp evidence score < 0.5). The resulting associations were manually curated to retain blood-related traits and hematological disorders based on hematological-associated keywords and domain expertise. Trait names were consolidated for interpretation.

Traits-gene association matrices were constructed separately for each PIP by pivoting L2G credible set scores (Open Targets) with traits as rows and genes as columns; missing values were set to zero. Heatmaps were generated using `clustermap` from Seaborn, with hierarchical clustering (Euclidean distance, average linkage) applied to genes (columns) on unscaled L2G scores. Traits (rows) were not clustered and were grouped according to predefined lineage categories. No row or column scaling was applied prior to clustering. Color scales represent L2G scores (vmin = 0.25). Traits were annotated by lineage category using a row color bar.

Using the subset of traits supported by curated genetic disorder evidence (ClinVar or PanelApp), traits were manually grouped into seven related hematological disorder categories (Mixed/Haematopoietic, Red Cell, MK/Platelet, Myeloid, Lymphoid, Immunodeficiency and Systemic/Developmental). For each PIP, the number of unique traits per category was counted and converted to within-PIP proportions. PIPs were hierarchically clustered using the same distance metric and linkage strategy computed on category-proportion vectors and were visualized as 100% stacked horizontal bar plots, with total trait counts annotated. Enrichment of disease gene categories within PerturbGen-defined clusters was assessed using one-sided Fisher’s exact tests. For each cluster–disease pair, a 2×2 contingency table was constructed comparing the number of disease-associated genes within the cluster to those outside the cluster, using the full set of perturbation-profiled genes as the background. Odds ratios were derived from the corresponding contingency tables, and *P* values were computed using the hypergeometric distribution as implemented in SciPy. Multiple testing correction across all cluster–disease comparisons was performed using the Benjamini–Hochberg procedure to control the false discovery rate (FDR).

### Pseudotime analysis

Pseudotime and lineage inference were performed using Moscot^40^ and Palantir^150^. A forward transition matrix was computed using the RealTimeKernel implemented in Moscot with self-transitions enabled, connectivity weighting (conn_weight = 0.2), and automatic thresholding. The transition matrix was imported into CellRank as a precomputed kernel and coarse-grained using GPCCA. Six macrostates were inferred, and dermal papilla (DP) cells were specified as terminal states. Fate probabilities toward the DPs were calculated for all cells.

Diffusion maps were computed using Palantir to model continuous pseudotime trajectories. A multiscale diffusion space was constructed for pseudotime ordering. Starting and terminal cells were defined based on annotated cell identities, with *FRZB*⁺ fibroblasts specified as early states and DP cells as terminal states. MAGIC^151^ imputation was applied for visualization of gene expression trends along trajectories with the default parameters.

### Sinkhorn divergence to assess transcriptomic similarity between skin organoid and fetal skin

Transcriptional similarity between organoid stromal populations and human fetal skin was quantified using entropic optimal transport distances (Sinkhorn divergence^99^). Organoid and fetal datasets were restricted to shared stromal cell types based on curated annotations (*FRZB*⁺ fibroblasts, *HOXC5*⁺ fibroblasts, *WNT2*⁺ fibroblasts, dermal condensate, dermal papilla and pre-dermal condensate); all other cell types were excluded.

To control for differences in cell number and cell type composition, stratified fixed-size subsampling was performed. For each organoid developmental stage and condition (perturbed or unperturbed), up to 1,000 cells were sampled while approximately preserving cell-type proportions and enforcing a minimum of 10 cells per cell type where possible. Fetal samples were subsampled per PCW to matched sizes.

Sinkhorn divergence was computed using the geomloss library (quadratic cost, ε = 0.05, scaling = 0.9, debiased formulation). For each organoid stage and fetal PCW, we calculated the Sinkhorn divergence between fetal samples and unperturbed organoids (d_unperturbed) and between fetal samples and perturbed organoids (d_perturbed). The difference in distances was defined as Δ = d_unperturbed − d_perturbed. Positive Δ values indicate reduced transcriptional distance of perturbed organoids to fetal tissue relative to unperturbed controls.

To account for variability introduced by subsampling, the full procedure was repeated 50 times with independent random seeds. Mean distances, standard deviations and empirical 95% confidence intervals were computed across repeats. Effect sizes were estimated using Cohen’s d.

