## Supplementary Figures for "Predicting how perturbations reshape cellular trajectories with PerturbGen"

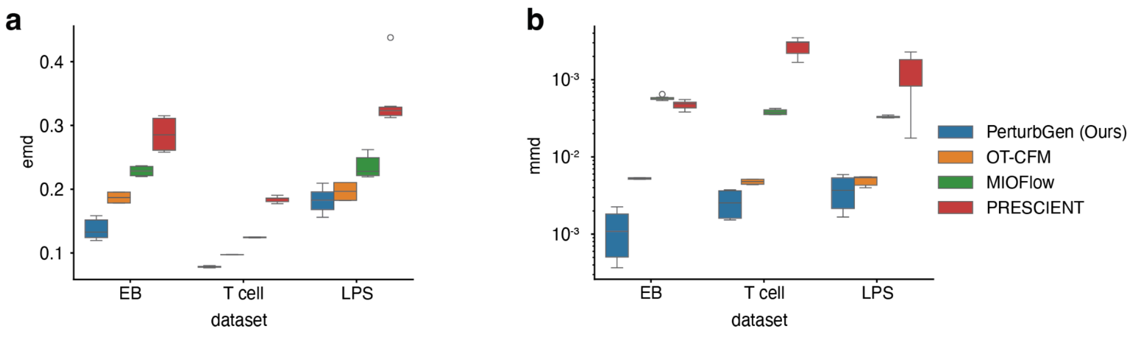


**Supplementary Fig. 1 | Single-cell time point benchmarking evaluating generative model performance.**

**a,** Prediction accuracy for held-out time points, quantified by Earth Mover's Distance (EMD), across the embryoid body, T cell and lipopolysaccharide datasets. **b,** Prediction accuracy quantified by Maximum Mean Discrepancy (MMD) on the same datasets; the y-axis is shown on a $\log_{10}$ scale. Results are averaged across held-out intermediate (interpolation) and subsequent (extrapolation) time points and across three independent random seeds. Center lines indicate the median, boxes the interquartile range and whiskers the full range. Methods are color-coded as follows: PerturbGen (blue), OT-CFM (orange), MIOFlow (green) and PRESCIENT (red). Lower values indicate improved predictive performance.

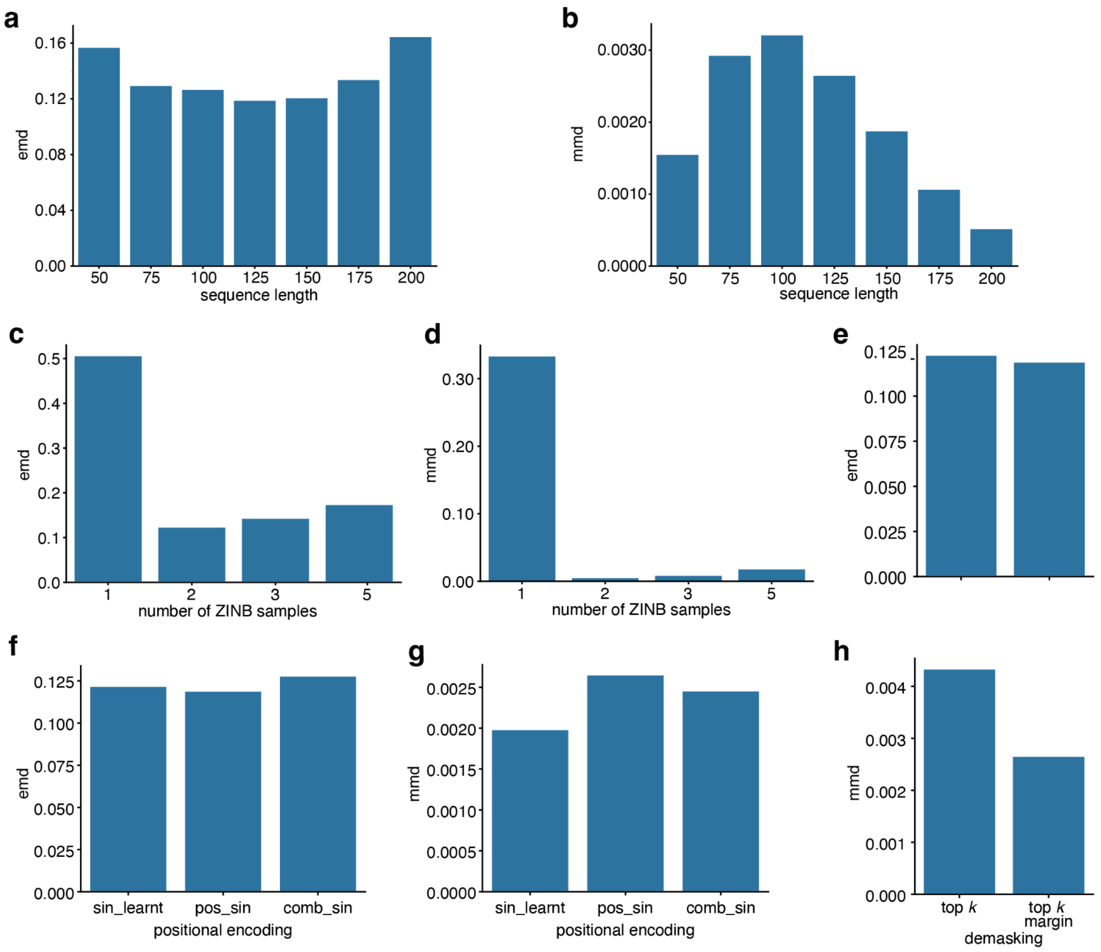


**Supplementary Fig. 2 | Single-cell time point benchmarking model ablation.**

**a,b,** Sequence-length ablation evaluated by Earth Mover's Distance (EMD) and Maximum Mean Discrepancy (MMD). **c,d,** Effect of zero-inflated negative binomial (ZINB) sampling on count prediction performance, quantified by EMD and MMD. **e,f,** Comparison of positional encoding strategies assessed by EMD and MMD. **g,h,** Iterative demasking strategy comparison evaluated by EMD and MMD.

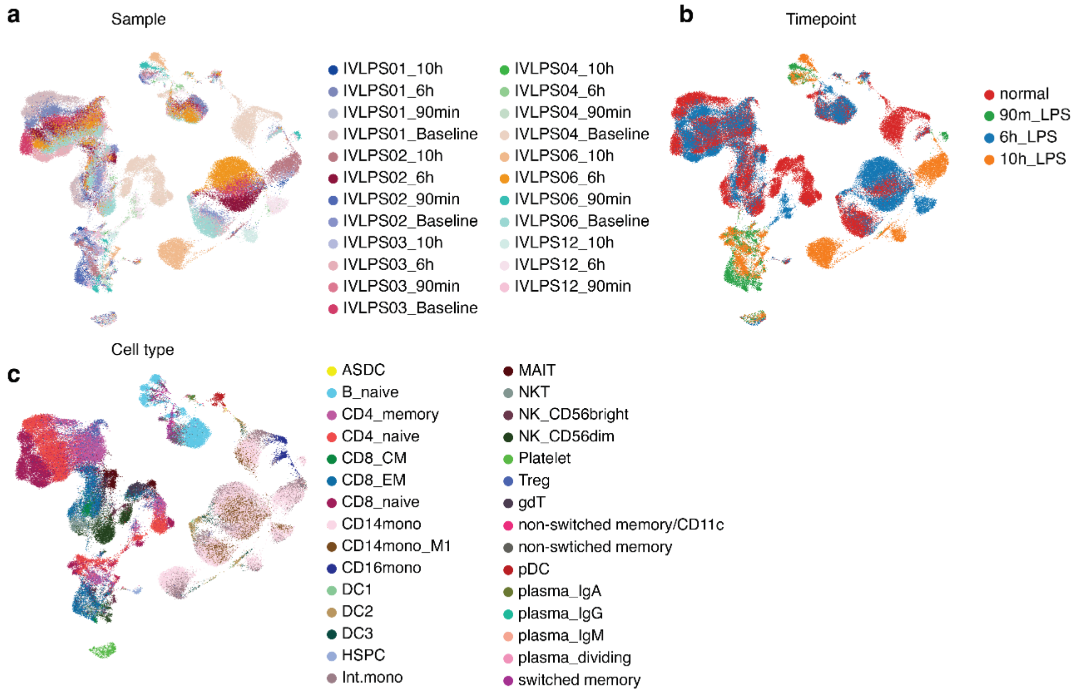


**Supplementary Fig. 3 | ScRNA-seq analysis of human PBMCs following *in vivo* LPS challenge.**

**a,** UMAP embedding of 23 samples included in the study. **b,** UMAP embedding colored by time point: baseline (normal), 90 min, 6 h and 10 h post-LPS infusion. **c,** UMAP embedding colored by cell type annotation.

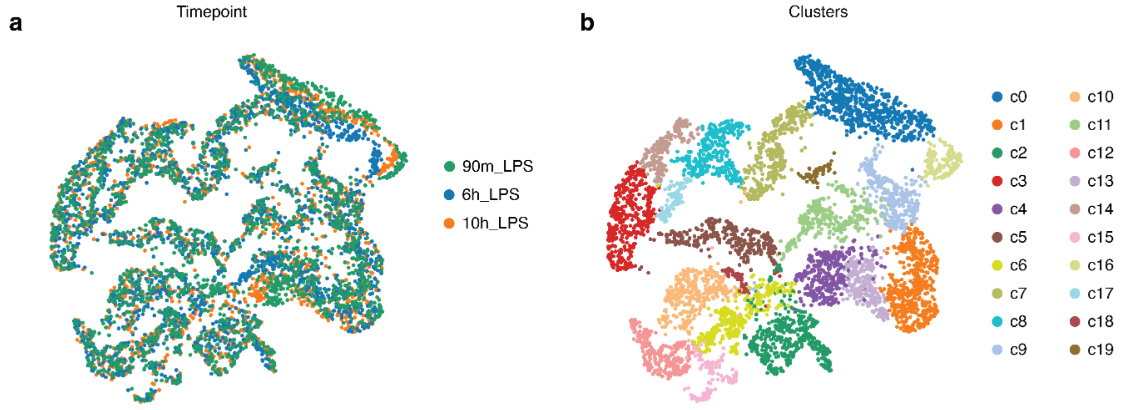


**Supplementary Fig. 4 | Time-resolved gene embedding analysis.**

**a,** UMAP embedding of gene embeddings colored by time point; each dot represents a single gene. **b,** UMAP embedding colored by Leiden clusters, which are defined as gene programs.

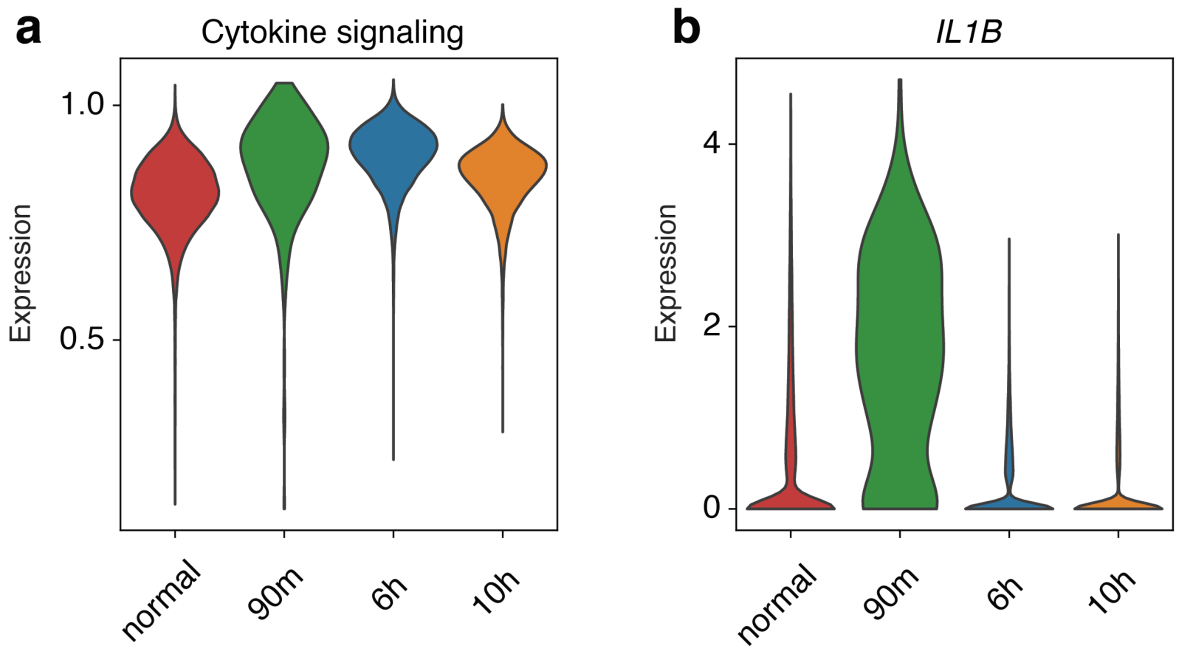


**Supplementary Fig. 5 | Cytokine signaling program and *IL1B* expression in myeloid lineages.**

**a,** Violin plots of cytokine signaling program across myeloid cell types at each time point. **b,** Violin plots of *IL1B* expression across myeloid lineages at each time point.

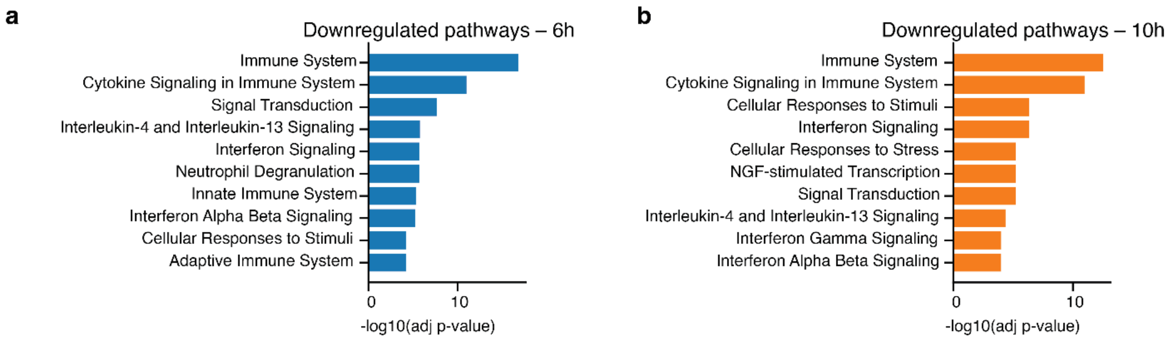


**Supplementary Fig. 6 | Over-representation analysis (ORA) of enriched pathways in the myeloid lineage following *in silico IL1B* knockout (KO).**
**a,b,** Bar plots showing the top 10 enriched pathways among downregulated differentially expressed genes (DEGs), ranked by $-\log_{10}$(adjusted *P* value), at 6 h **(a)** and 10 h **(b)**. DEGs were defined using an adjusted *P* value < 0.05 and $|\log_2$ fold change$| \geq 0.2$.

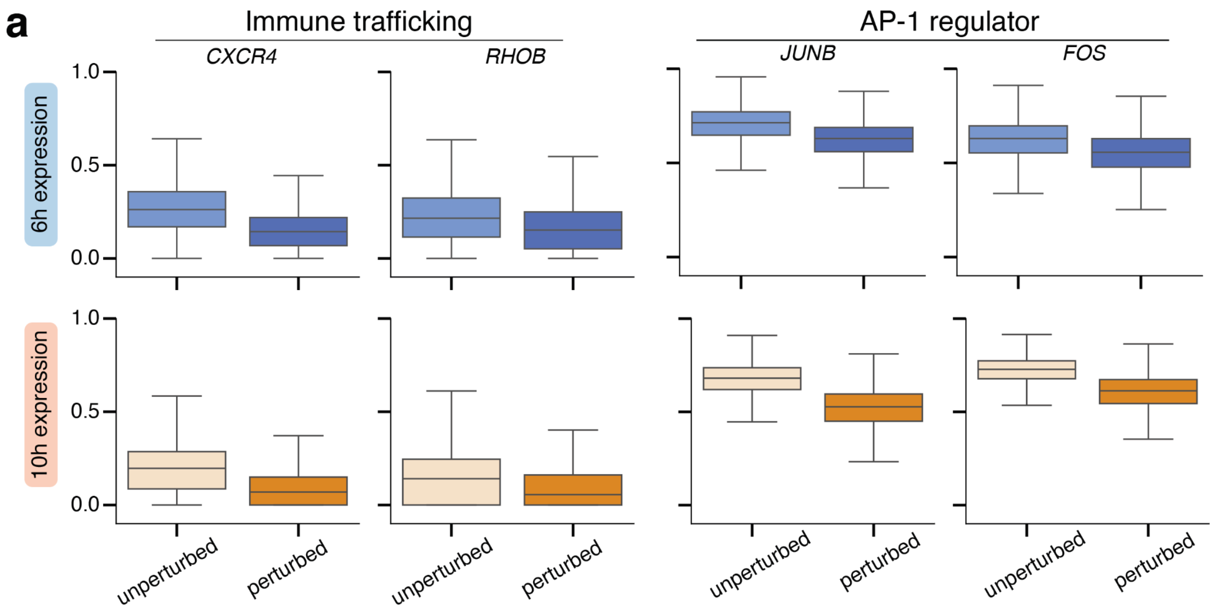


**Supplementary Fig. 7 | Representative genes associated with *IL1B* KO.**
**a,** Representative immune trafficking and AP-1 regulator genes showing lower predicted expression in perturbed compared with unperturbed cells at 6 h and 10 h. Box plots display predicted min–max normalized expression in unperturbed and perturbed cells at 6 h (top) and 10 h (bottom). Center lines indicate the median, boxes the interquartile range and whiskers the full range.

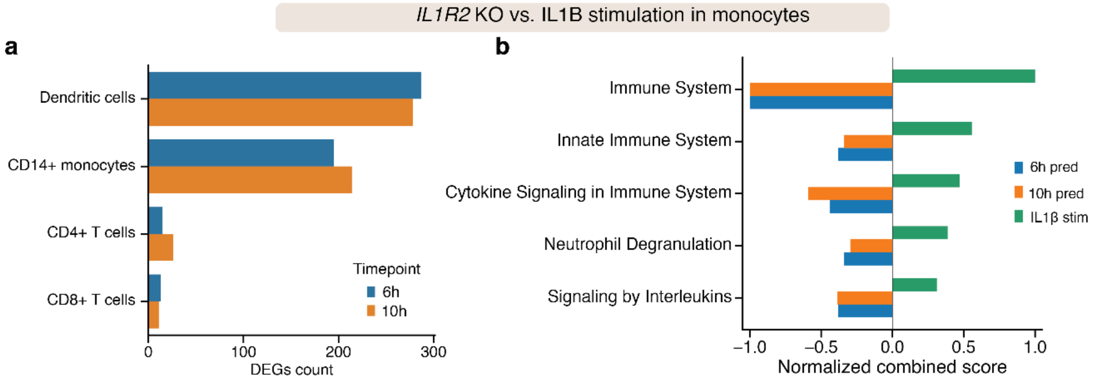


**Supplementary Fig. 8 | Quantitative and qualitative assessment of *in silico IL1R2* KO.**

**a,** Bar plot showing the perturbation effect quantified by the number of DEGs following *in silico IL1R2* KO. **b,** Mirrored bar plot showing the top five pathways ranked by combined score, illustrating directional effects relative to IL1β stimulation.

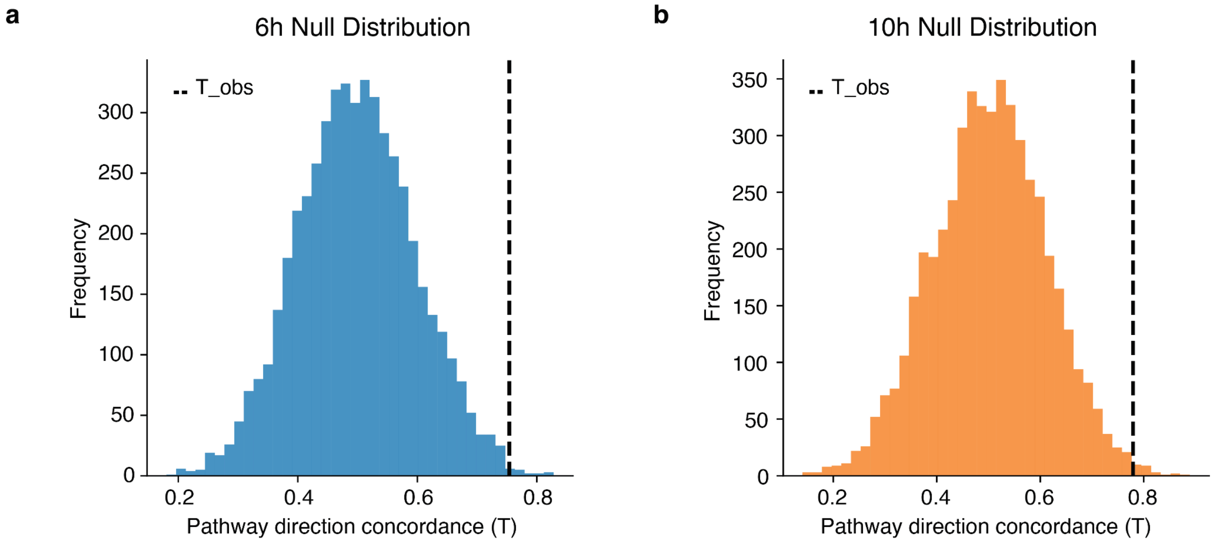


**Supplementary Fig. 9 | Permutation test assessing pathway-level reversal concordance between *in silico IL1R2 KO* and IL1β stimulation signatures.**

**a,b**, Null distributions of balanced accuracy for ORA-based pathway directionality prediction, generated by random permutation of gene labels prior to enrichment analysis using Reactome pathways at 6 h **(a)** and 10 h **(b)**. Dashed vertical lines indicate the observed model performance relative to the empirical null distribution.

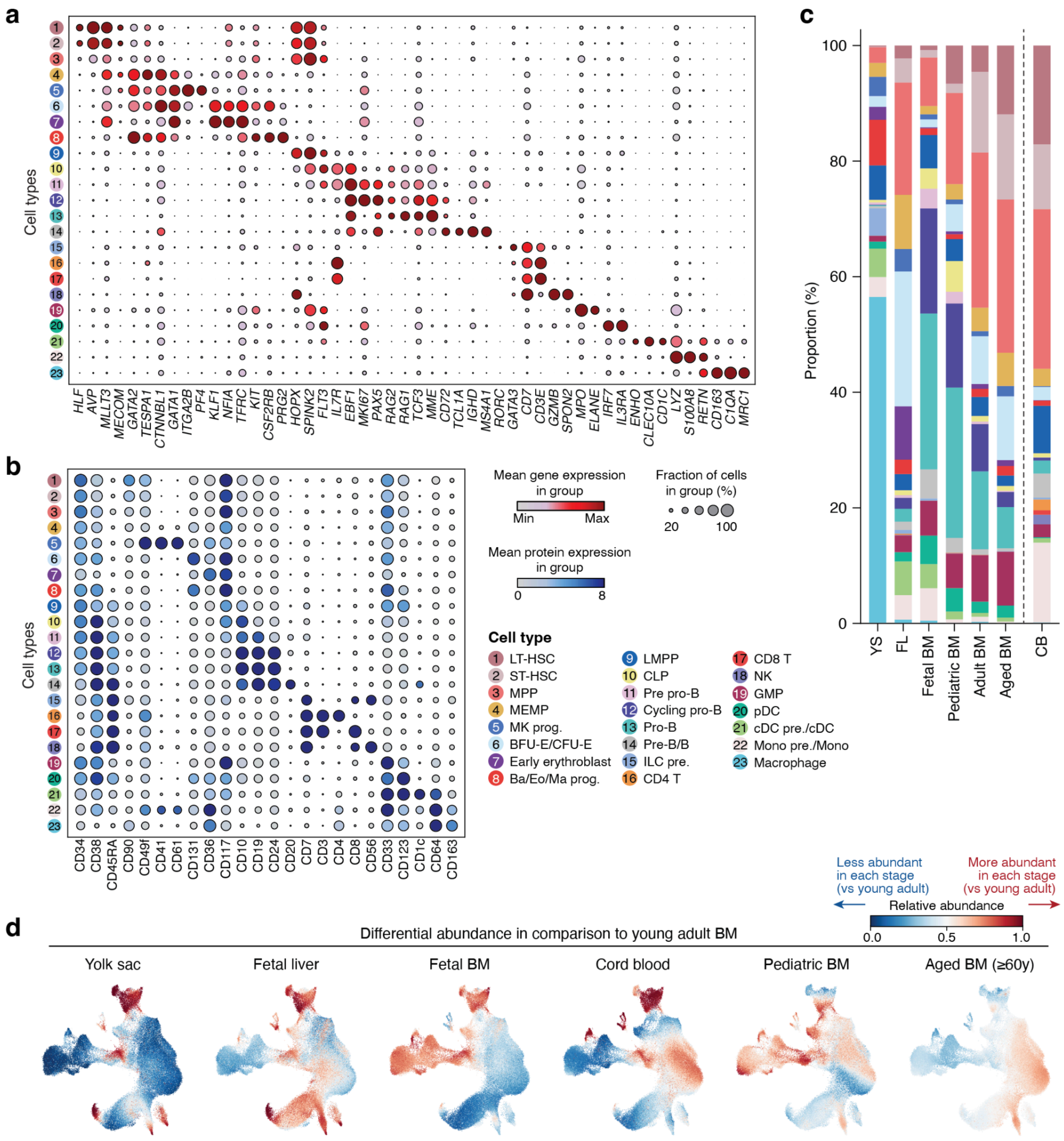


**Supplementary Fig. 10 | Hematopoietic stem and progenitor cells across the human lifespan.**

**a,b,** Dot plots showing the cluster-mean expression of cell type marker genes (**a**) and proteins (**b**). **c**, Hematopoietic cell type composition by developmental stage. **d**, Differential abundance landscapes of age-specific hematopoiesis. Young adult bone marrow samples (29–50 years old) were used as the reference group and compared against each age group.

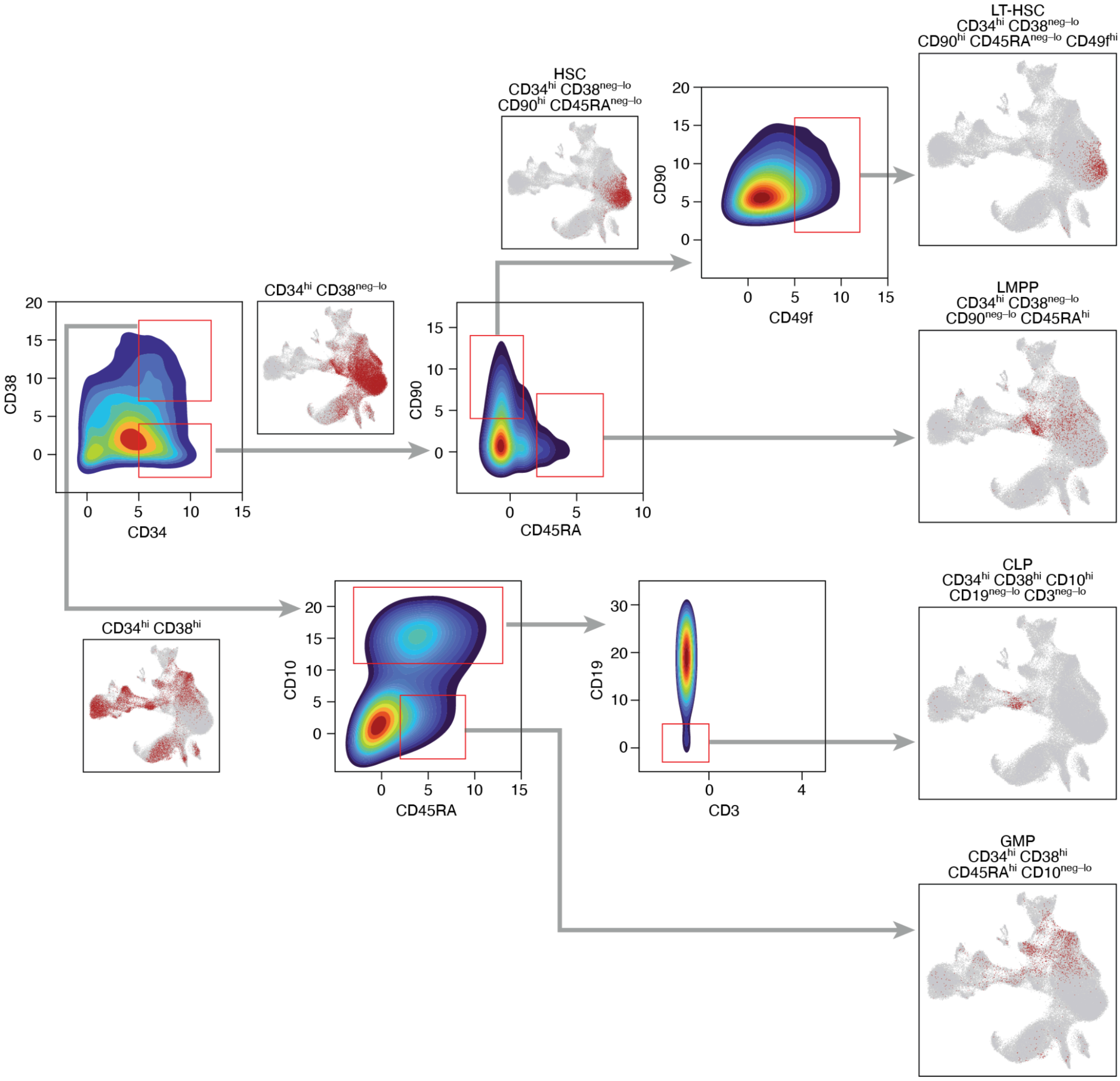


**Supplementary Fig. 11 | HSPC gating strategy based on protein markers for cell type annotation.**

*In silico* gating approach for the identification of distinct immunophenotypic populations. Two-dimensional density plots show denoised protein expression. Isolated cells are depicted in red in the UMAP plots.

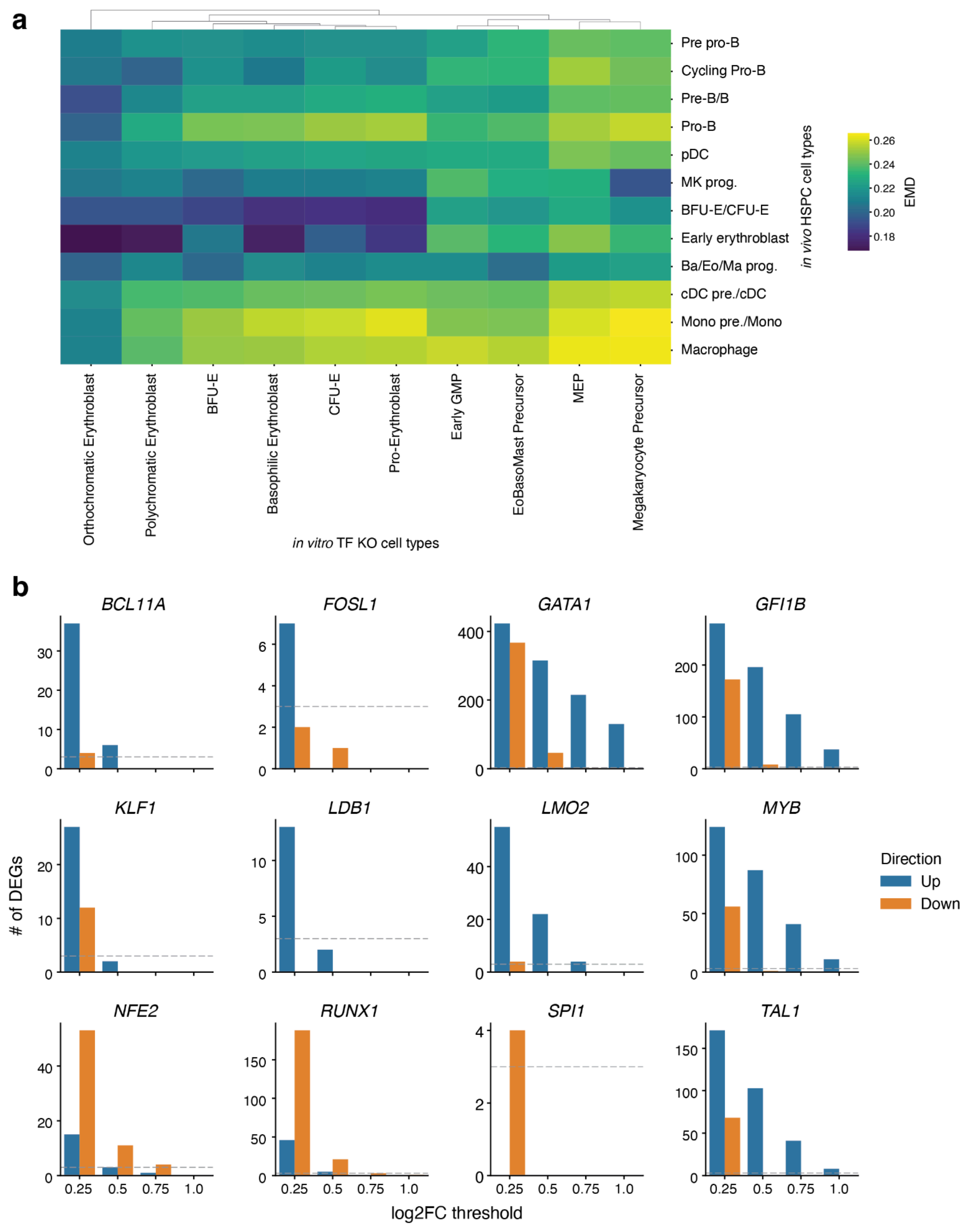


**Supplementary Fig. 12 | Criteria for cross-dataset cell type alignment and perturbation selection**

**a,** Cross-dataset cell type similarity quantified using Earth Mover's Distance (EMD; lower values indicate greater similarity) between the *in vitro* CRISPR transcription factor (TF) knockout (KO) dataset and the *in vivo* HSPC dataset. Based on EMD ≤ 0.2, erythroid and megakaryocytic lineage cell types were retained for downstream analyses, while early GMP, MEP, and EoBasoMast precursor populations were excluded. **B,** Number of up- and downregulated genes identified for twelve TF KOs with significant transcriptional effects (adjusted $P$ value < 0.05). The dotted line indicates the minimum DEG threshold (≥ 3 genes) required for the perturbation benchmark.

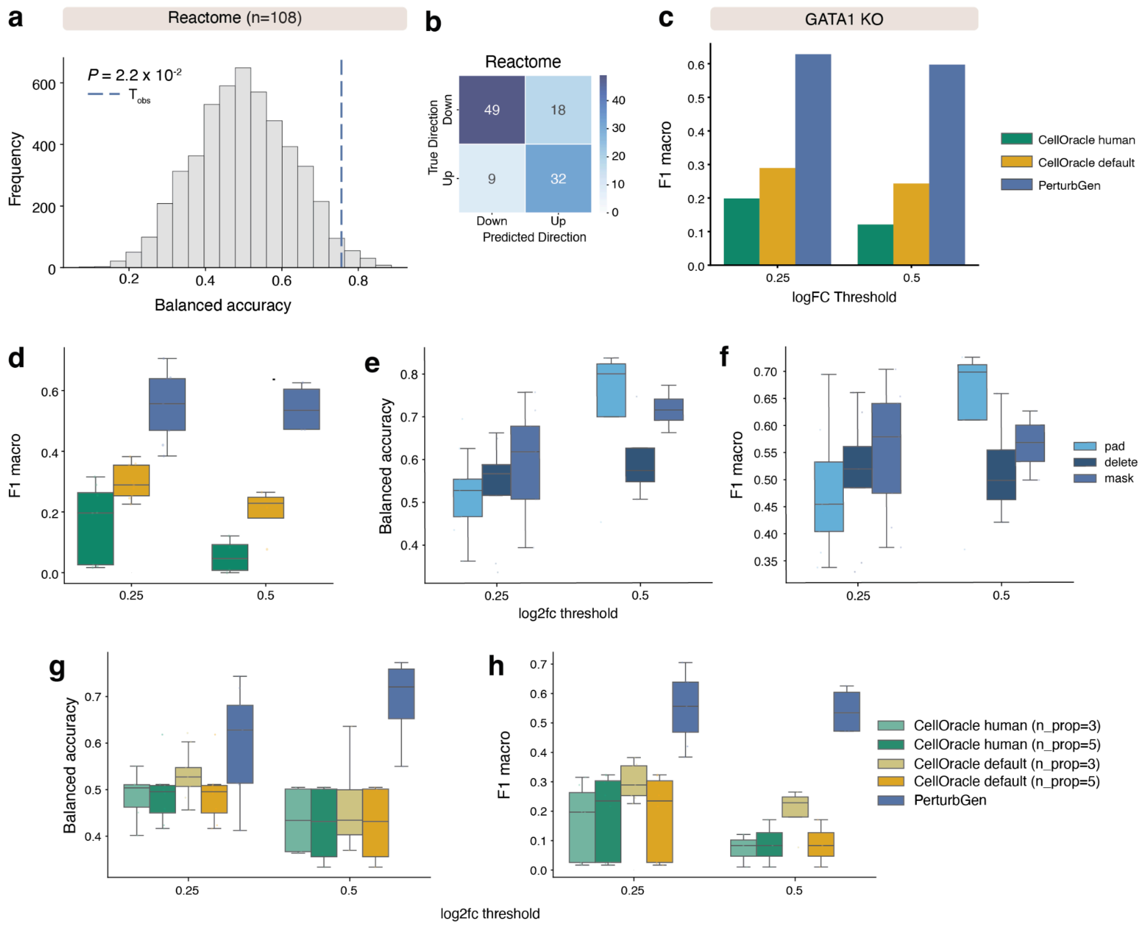


**Supplementary Fig. 13 | Perturbation benchmarking comparing PerturbGen predictions to *in vitro* CRISPR transcription factor knockout data.**

**a,** Pathway-level concordance between in silico and in vitro GATA1 knockout, quantified using normalized enrichment scores (NES) from gene set enrichment analysis (GSEA) of Reactome pathways (n = 108 gene sets with FDR < 0.05 in the *in vitro* dataset). Model performance was evaluated using balanced accuracy for pathway directionality (up- versus down-regulated) and compared to a permutation-based null distribution (empirical $P$ value shown). Empirical $P$ values were estimated from 5,000 permutations (see Methods). The dashed vertical line indicates observed model performance. **b,** Confusion matrix summarizing pathway comparing predicted with experimentally observed pathway directionality for Reactome pathways. **c, d,** Benchmarking of GATA1 KO (c) and nine transcription factors (d), PerturbGen with CellOracle using either the default gene regulatory network (GRN) or a human ATAC-derived GRN. Performance is quantified by F1 macro score across $\log_2$ fold-change thresholds. **e,f,** Model ablation analysis evaluating strategies to introduce perturbation-padding, deletion, or masking of the perturbed gene-quantified by balanced accuracy (e) and F1 macro score (f). Metrics equally weight up- and down-regulated classes. **g,h,** Benchmarking comparison with CellOracle using varying numbers of propagation steps (3 or 5). Performance is quantified by balanced accuracy (g) and F1 macro score (h) across $\log_2$ fold-change thresholds. Boxes indicate the interquartile range (IQR), center lines the median, whiskers extend to 1.5× IQR.

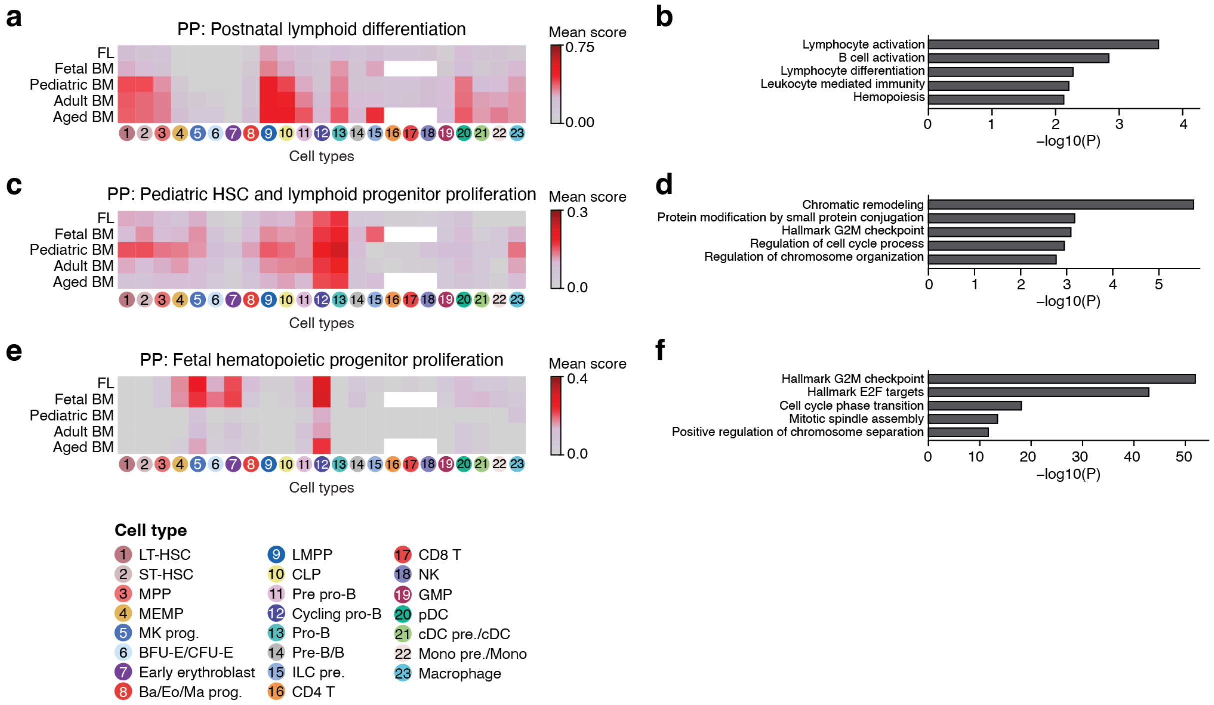


**Supplementary Fig. 14 | Perturbation-induced program (PIP) annotation through gene scoring and GSEA.**

**a-f,** Gene score heatmaps (**a**,**c**,**e**) and enriched gene ontology terms (**b**,**d**,**f**) for the postnatal lymphoid differentiation program (**a**,**b**), pediatric HSC and lymphoid progenitor proliferation program (**c**,**d**), and fetal hematopoietic progenitor proliferation program (**e**,**f**). In the heatmaps (**a**,**c**,**e**), cluster-mean gene scores in each developmental stage are plotted.

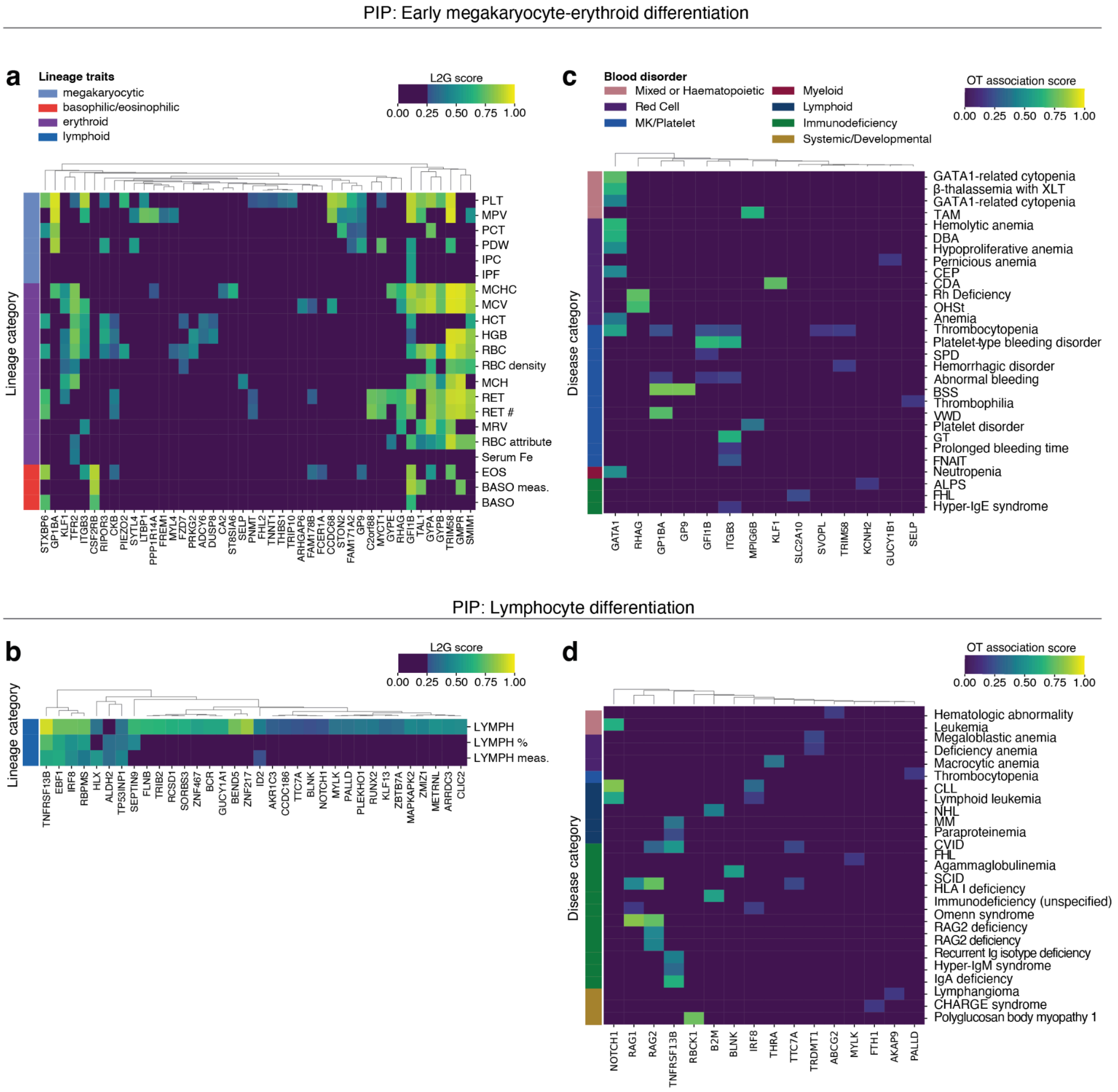


**Supplementary Fig. 15 | Mapping PIPs to hematological traits and blood disorders.**

**a,b,** Hierarchically clustered heatmaps showing associations between genes assigned to the early megakaryocyte–erythroid differentiation (a) and lymphocyte differentiation (b) PIPs and hematological traits from GWAS. Associations are colored by Open Targets Locus-to-Gene (L2G) scores (≥0.25). Row annotations indicate lineage categories for each trait. **c,d,** Hierarchically clustered heatmaps showing associations between genes assigned to the early megakaryocyte–erythroid differentiation (c) and lymphocyte differentiation (d) PIPs and blood disorders. Associations are colored by weighted OT association scores. Row annotations indicate disease category.

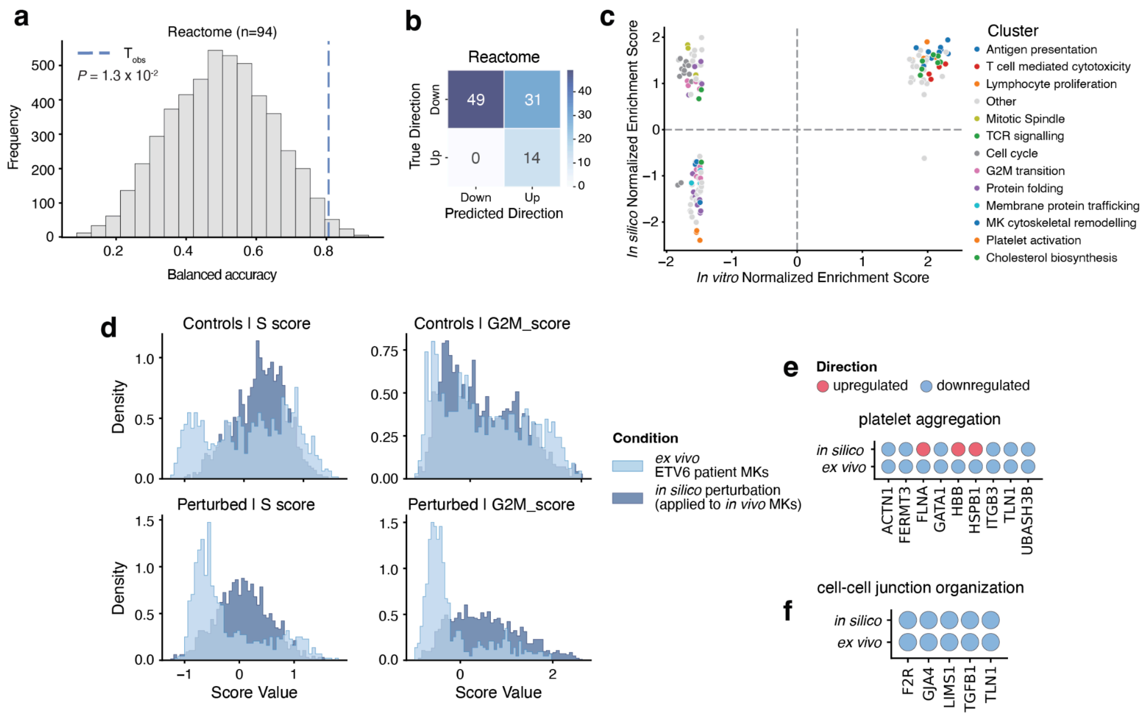


**Supplementary Fig. 16 | *In silico ETV6* perturbation recapitulates disease-associated transcriptional programs observed *ex vivo***

**a,** Pathway-level concordance between *in silico ETV6* perturbation *and ex vivo* samples from patients with *ETV6*-associated thrombocytopenia, quantified using NES from GSEA of Reactome pathways (n = 94 gene sets with FDR < 0.05 in the *ex vivo* dataset). Model performance was evaluated using balanced accuracy for pathway directionality (up- versus down-regulated) and compared to a permutation-based null distribution (empirical *P* value shown). Empirical *P* values were estimated from 5,000 permutations (see Methods). The dashed vertical line indicates observed model performance. **b,** Confusion matrix summarizing pathway comparing predicted with experimentally observed pathway directionality for Reactome pathways. **c,** Comparison of NES between *in silico* and *ex vivo* conditions; each point represents a gene set. Gene sets are colored by hierarchical clustering and manually annotated functional categories. **d,** Distribution of cell cycle phase scores (S phase and G2M scores), shown separately for baseline control and perturbed conditions in *in vivo* and *ex vivo* datasets. **e,f,** Concordance in directionality of GSEA leading edge genes for platelet aggregation (e) and cell-cell junction organization (f) between *in silico* and *ex vivo* conditions. Dots are colored by $\log_2$ fold-change sign (red, positive; blue, negative).

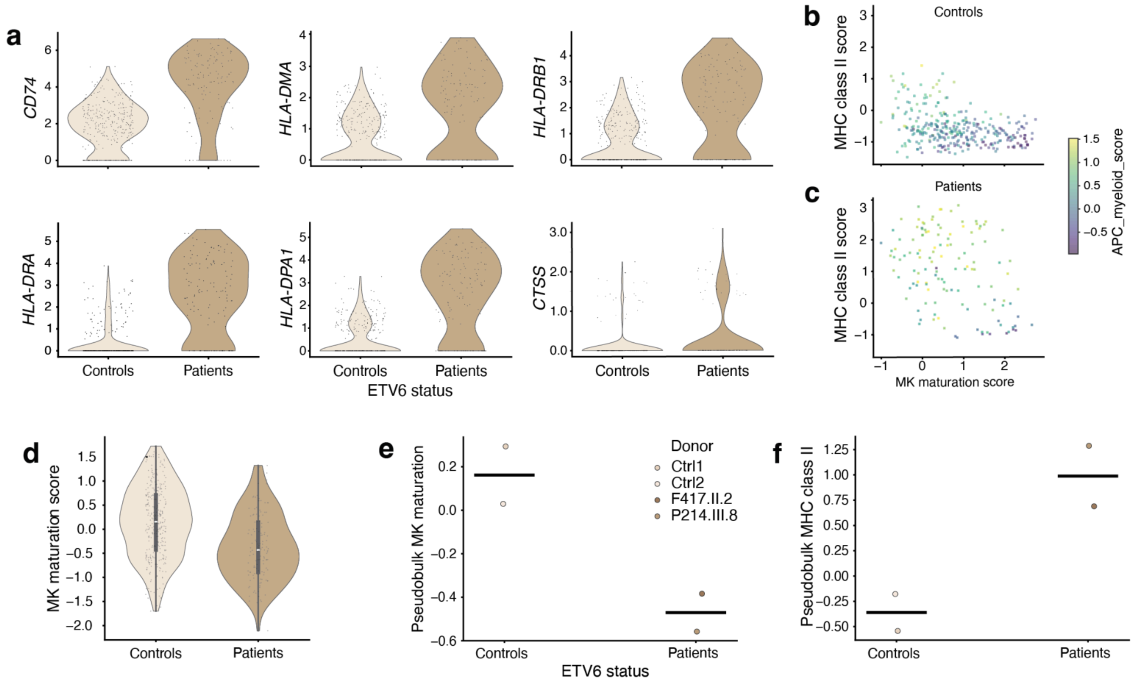


**Supplementary Fig. 17 | Altered MHC class II and maturation programs in megakaryocytes in ETV6-associated thrombocytopenia**

**a,** Expression levels for MHC class II-associated genes (*CD74*, *HLA-DRA*, *HLA-DMA*, *HLA-DPA1*, *HLA-DRB1* and *CTSS*) in megakaryocyte (MK) marker-positive and antigen-presenting cell (APC) marker-negative cells, stratified by patient condition (Methods). **b,c,** Scatter plots of MK maturation score versus MHC class II score, colored by APC myeloid score, shown for healthy controls (b) and patients (c). **d,** Distribution of MK maturation scores stratified by condition. **e,f,** Pseudobulk expression values for MK maturation score (e) and MHC class II score (f), aggregated per donor across conditions.

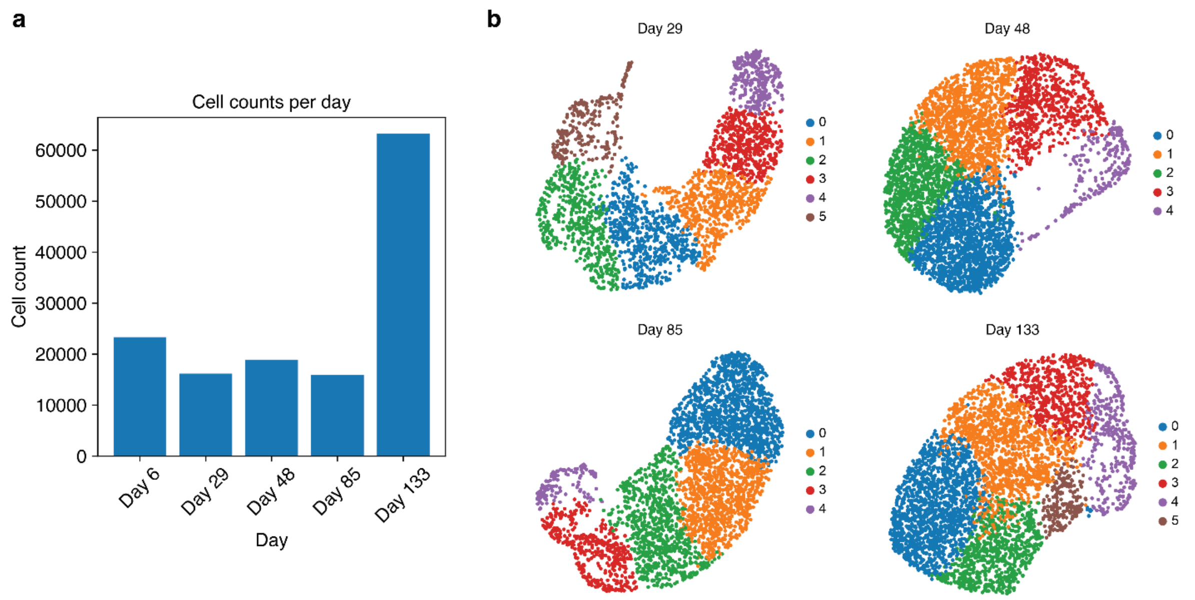


**Supplementary Fig. 18 | Large-scale time-resolved perturbation atlases in a published human skin organoid dataset.**

**a,** Number of skin organoid cells across cultured time points. **b,** UMAP embedding of perturbation atlases predictions across culture time points; each dot represents a single gene perturbation, mean-pooled across cells with detectable expression of the perturbed gene.

**a**

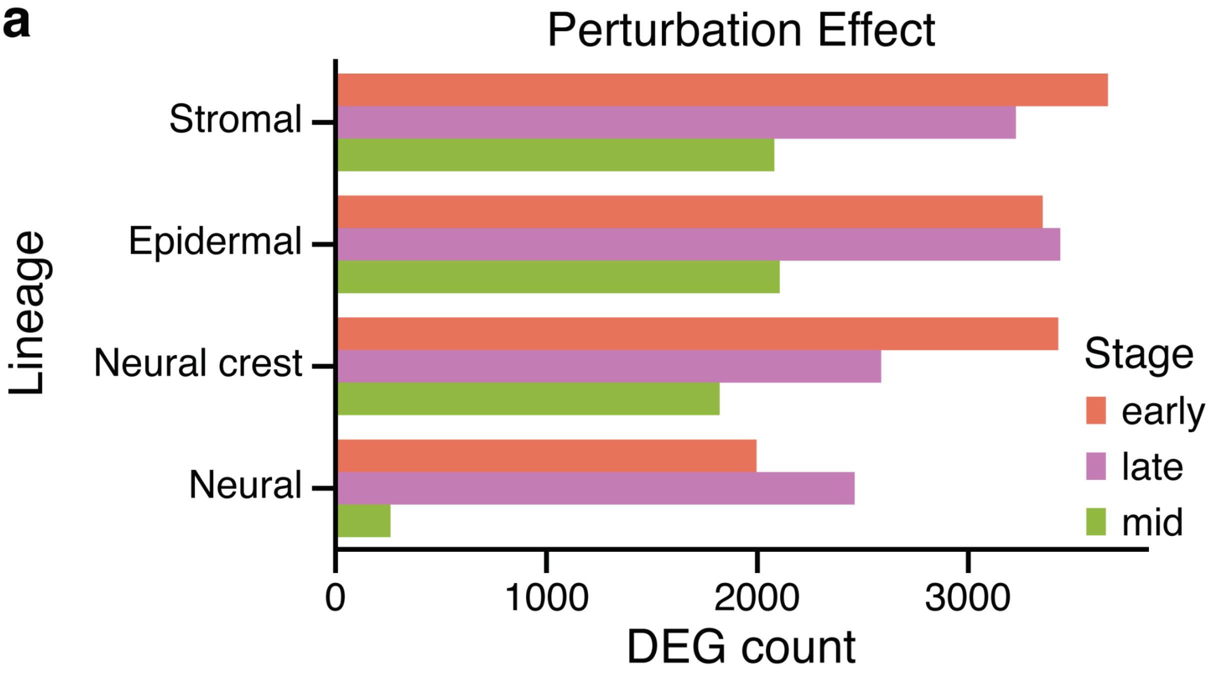


**Supplementary Fig. 19 | Lineage-aware perturbation effect of *in silico GSK3B* KO in human skin organoids.**

**a,** Bar plot of perturbation effect quantified by the number of significant DEGs (adjusted *P* value < 0.05) across lineages stratified by developmental stages.

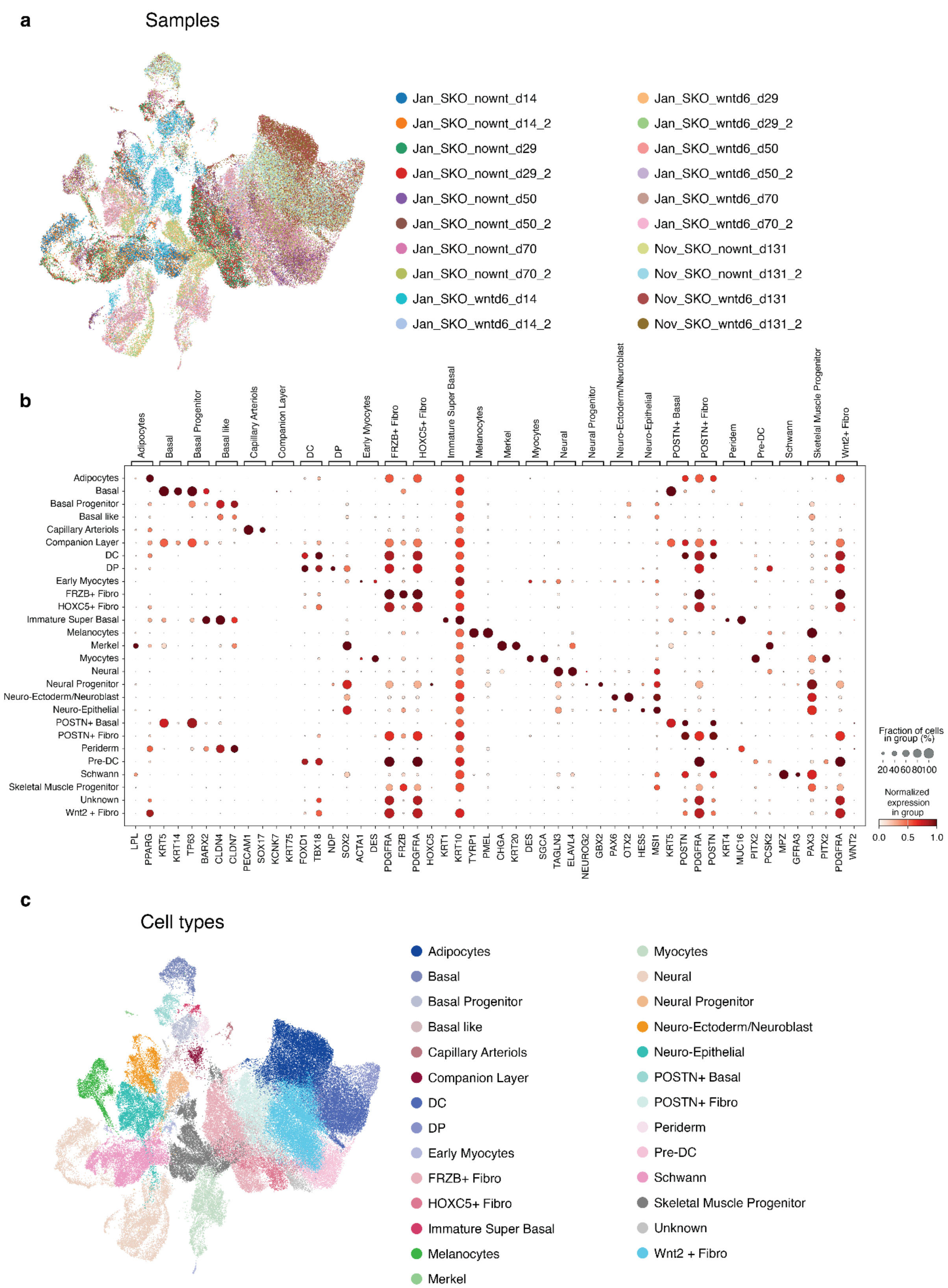


**Supplementary Fig. 20 | ScRNA-seq profiling of human skin organoids following *in vitro* GSK3β inhibition.**

**a,** UMAP embedding of all 20 samples with 2 biological replicates per each condition colored by sample identity. **b,** Dot plot of normalized expression of canonical marker genes used for cell type annotations. **c,** UMAP embedding colored by cell type annotations.

**a**

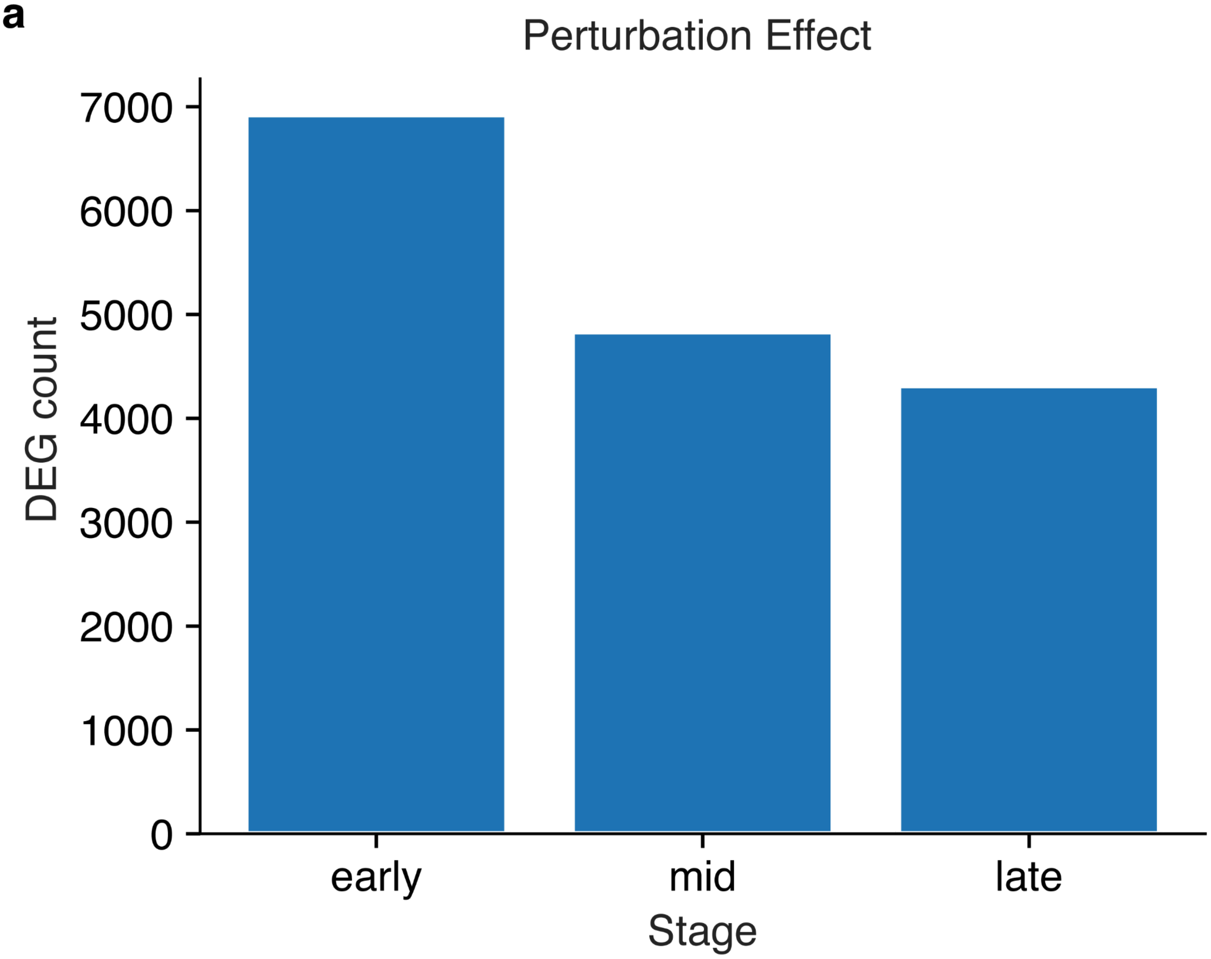


**Supplementary Fig. 21 | *In vitro* GSK3β inhibition effect in skin organoid stromal development.**

**a,** Bar plot showing the perturbation effect following *in vitro* GSK3β inhibition, quantified by the number of DEGs across developmental stages.

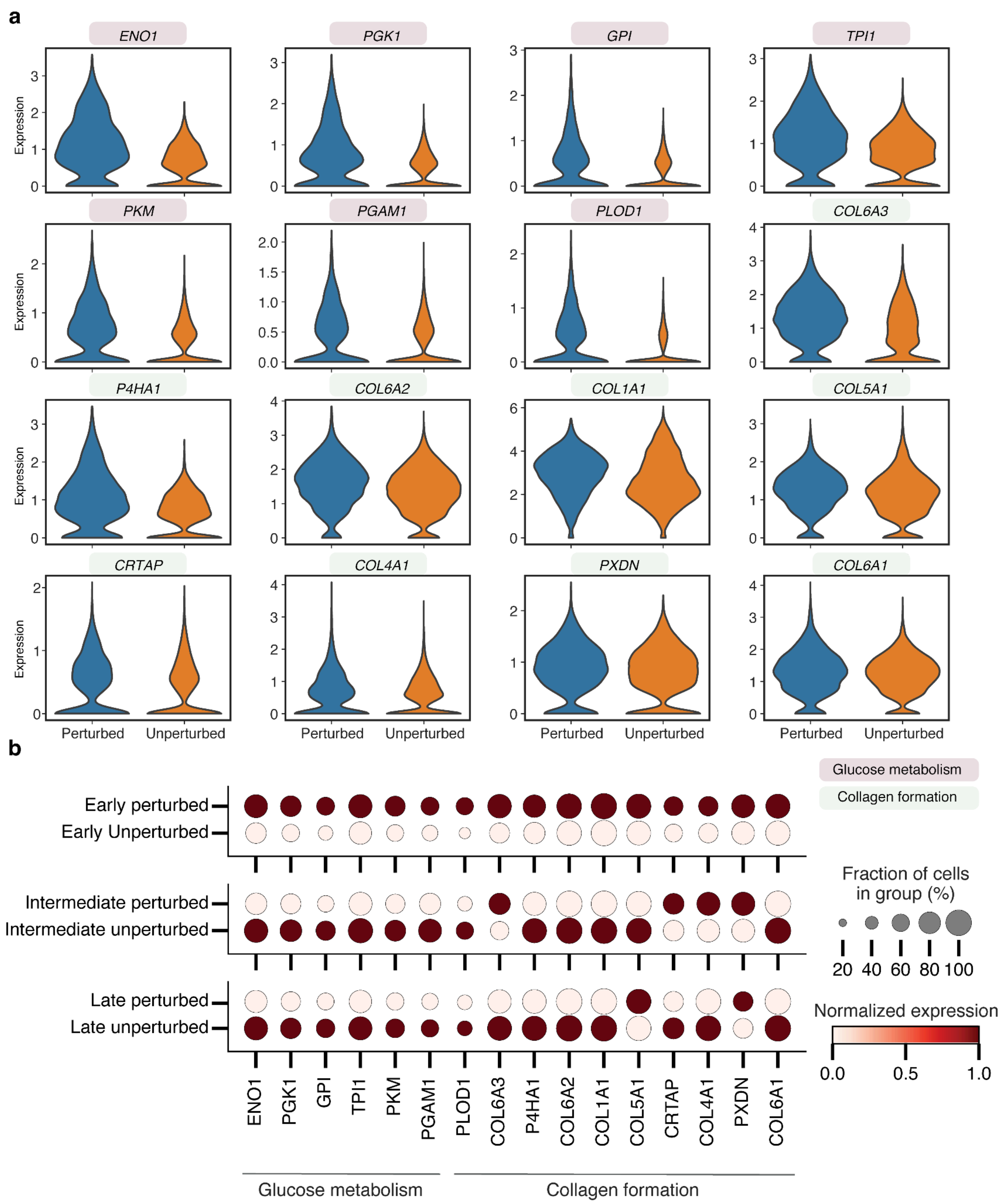


**Supplementary Fig. 22 | Gene expression changes in glucose metabolism and collagen-related genes following *in vitro* GSK3β inhibition.**

**a,** Violin plots showing the mean expression values of representative genes involved in glucose metabolism and collagen formation in perturbed (blue) and unperturbed (orange) stromal cells of skin organoids. **b,** dotplots show the normalized expression values of representative genes from mentioned pathways across different stages of organoid development.

a

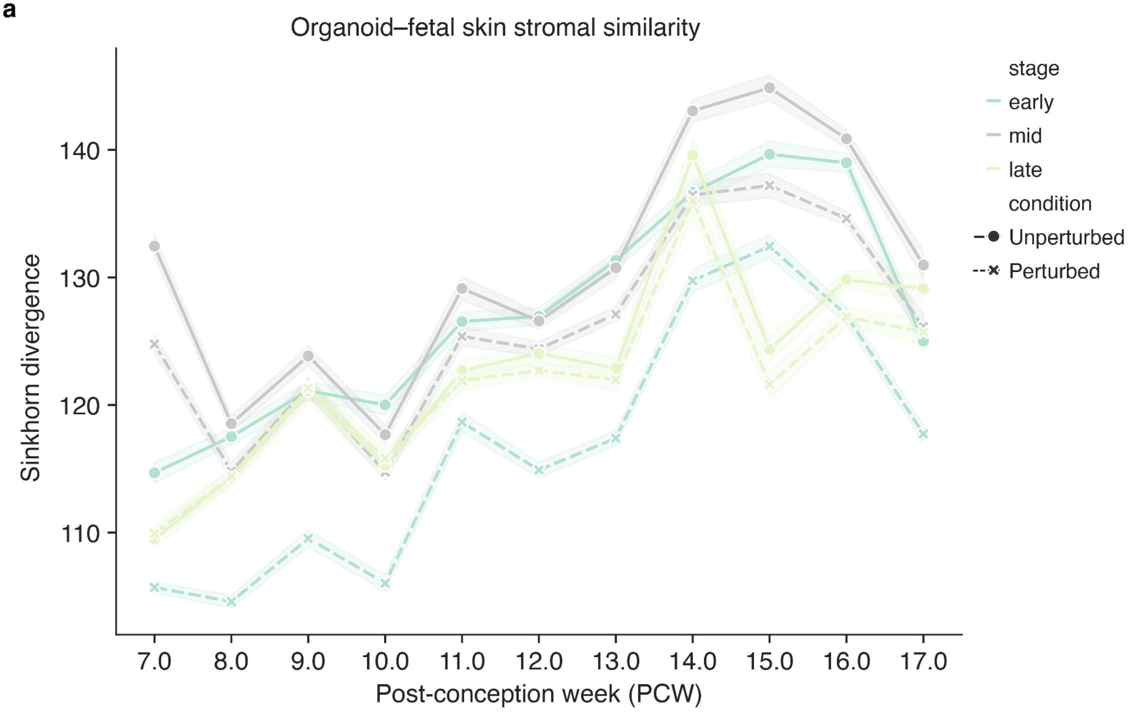


**Supplementary Fig. 23 | Quantitative assessment of similarity between GSK3β-inhibited stromal in skin organoids and fetal skin stromal.**

**A,** Sinkhorn divergence between organoid stromal and human fetal skin stromal across post-conceptional weeks (PCWs). For each PCW and condition, values were measured across organoid stromal stages, with error bands representing the standard deviation of stage-specific distances.
